# Beyond dimension reduction: Stable electric fields emerge from and allow representational drift

**DOI:** 10.1101/2021.08.22.457247

**Authors:** Dimitris A. Pinotsis, Earl K. Miller

**Author notes:** **Correspondence**: Dimitris A. Pinotsis, Centre for Mathematical Neuroscience and Psychology and Department of Psychology, City —University of London, London EC1V 0HB, United Kingdom.

## Abstract

It is known that the exact neurons maintaining a given memory (the neural ensemble) change from trial to trial. This raises the question of how the brain achieves stability in the face of this representational drift. Here, we demonstrate that this stability emerges at the level of the electric fields that arise from neural activity. We show that electric fields carry information about working memory content. The electric fields, in turn, can act as “guard rails” that funnel higher dimensional variable neural activity along stable lower dimensional routes. We obtained the latent space associated with each memory. We then confirmed the stability of the electric field by mapping the latent space to different cortical patches (that comprise a neural ensemble) and reconstructing information flow between patches. Stable electric fields can allow latent states to be transferred between brain areas, in accord with modern engram theory.

## Introduction

In the era of large scale electrophysiology ^1^, neural recordings of high dimensionality are abundant. Yet this has revealed that brain areas seems to exchange information in low dimensions, using few task-related variables (latent variables) ^2^. Indeed, brain dynamics evolve in low, not high, dimensional spaces ^3,4^. These spaces are found by dimensionality reduction, ^5,6^. Low dimensionality underlies a variety of cognitive and motor tasks ^7,8^.

A key point is that low-dimension latent variables track information and task demands and are stable, highly correlated across trials ^9^. This stands in contrast to higher-dimensional neural dynamics; while there is some overlap^10^, the specific neurons and synapses activated are variable across trials ^11,12,13^. This appears paradoxical: which specific neurons are activated continuously changes, synapses rewire etc., yet at the functional/behavioral, stability comes from low dimensional, latent variables ^14–16^. This low dimensional stability is important for normal cognition and behavior. Downstream neurons and networks need some consistency from upstream networks even though those upstream networks are under continuous reconfiguration.

The continuous reconfiguration is known as representational drift ^17^. It occurs at a time scale of days, minutes or seconds ^18^. It helps ensure the robustness of brain circuits. If some neurons fail, others can do the same task ^19^. Plus, neurons, especially in higher cortical areas, have mixed selectivity which adds computational horsepower and cognitive flexibility ^20,21^. Representational drift may also be important for the brain computations needed for Predictive Coding ^22^ and Reinforcement Learning ^23^. But the biophysical mechanism that allows low-dimensional brain dynamics to emerge despite the representational drift, is still a mystery.

Here, we suggest that this low dimensional stability is an emergent property of the electric fields generated by neural activity. Consider the following: First, that ensembles are functionally integrated within larger brain networks ^24–26^. Networks must somehow represent the same memory at different times even though larger networks in which they embedded are in different states at different times. Given this fluctuating network activity, it is difficult to imagine how that memory could be represented by a specific set of neurons and connections, even if one assumes redundancy. Second, different combinations of electric sources can generate the same field ^27^. Taken together, the above two facts suggest that a changing input from the rest of the brain leads to a reconfiguration of the ensemble so that a stable electric field is maintained. Thus, a stable electric field level emerges from a high-dimensional representational drift of specific neurons.

It may help to consider the following analogy: Brain anatomy is like the road-and-highway system. It is where traffic could go. Current thoughts, memories etc. are the patterns of traffic at that moment. An exact network of specific neurons is one particular route through the road- and-highway system. But, importantly, the same destination can be reached by taking different routes at different times (i.e., representational drift). What really matters are the general patterns of the traffic, *not* the exact roads it takes. There are multiple ways to travel from location A to location B.

This motivates the following hypothesis: That ensemble representation at the electric field level is more robust and less variable than representation at the level of specific neurons and circuits. If true, this could explain how low-dimensional stable computations arise despite representational drift.

Here, we tested whether this hypothesis is supported by data from a spatial delayed saccade task. We characterized the stability of both the electric field and of the neural activity that generates this field. The same data were earlier used to build brain computer interfaces ^28^ and provide an neurobiological explanation of the oblique effect^29^. We here used them to train a biophysical neural network model as an autoencoder that learned to maintain spatial locations. This gave us the latent space similarly to other dimensionality reduction approaches ^8,30^. Then, we went one step further. We obtained single trial estimates of effective connectivity between different neurons. These describe how information propagates over a cortical patch occupied by the neural ensemble; and how neurons communicate via electric signals sent from one part of the patch to the other. This is a difference between our approach and other approaches. Our approach maps the latent space to a cortical patch. It goes beyond dimensionality reduction and reconstructs information flow.

Following ^29^, we reconstructed the effective connectivity between neurons on the patch from the latent space obtained earlier. These connectivity estimates describe the exchange of electric signals within the ensemble. This extra step also allowed us to reconstruct the electric field produced by the ensemble. Having a detailed description of electric signals and neural activity within the patch, we computed the electric field near it, using a classic dipole model from electromagnetism ^31^. To sum up, we predicted neural activity and the electric field generated each time (trial) the same location had to be remembered. Then, we tested if they were the same across trials. We found that the electric field was different for different remembered locations and highly consistent across trials. It also contained stable information about the remembered locations, while specific neurons activated were variable across trials (representational drift).

## Methods

### Experimental Data and Recording Setup

We reanalyzed data from ^28^. The same data were used in our earlier paper ^29^. Two adult male monkeys (monkey C, Macaca fascicularis, 9kg; monkey J, Macaca mulatta, 11kg) were handled in accordance with National Institutes of Health guidelines and the Massachusetts Institute of Technology Committee on Animal Care. They were trained to perform an oculomotor spatial delayed response task (Supplementary Figure 1B). This task required the monkeys to hold the location of one of six randomly chosen visual targets (at angles of 0, 60, 120, 180, 240 and 300 degrees, 12.5-degree eccentricity) in memory over a brief (750 ms) delay period and then saccade to the remembered location. If a saccade was made to the cued angle, the target was presented with a green highlight and a water reward was delivered otherwise the target was presented with a red highlight and reward was withheld. Three 32-electrode chronic arrays were implanted unilaterally in PFC, SEF and FEF in each monkey (Supplementary Figure 1C). Each array consisted of a 2 x 2 mm square grid, where the spacing between electrodes was 400 um. The implant channels were determined prior to surgery using structural magnetic resonance imaging and anatomical atlases. From each electrode, we acquired local field potentials (extracted with a fourth order Butterworth low-pass filter with a cut-off frequency of 500Hz, and recorded at 1 kHz) using a multichannel data acquisition system (Cerebus, Blackrock Microsystems). We analyzed local field potentials (LFPs) during the delay period when monkeys held the cued angles in memory.

### From the Wilson Cowan equations to Deep Neural Fields

Below we derive the evolution equations for a biophysical neural network model whose connectivity parameters have been obtained after training it as an autoencoder. This describes the activity of a neural ensemble. Its connectivity is such that the mutual information between the remembered cue and the ensemble activity is maximized. The corresponding weights are optimal in an information-theoretic sense.

Consider a neural ensemble that consists of neurons occupying a cortical patch (two dimensional Euclidean manifold) *M_A_*. Let *u_a_, v_a_* be two spatial variables parameterizing a *M_a_*, (*u_a_, v_a_*) ∈ *M_A_*, see e.g.^32–34^. Let 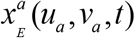 and 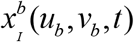 be the membrane potential of excitatory neurons and inhibitory neurons at locations (*u_a_, v_a_*) and (*u_b_, v_b_*) on the cortical surface and time *t*. The time evolution of 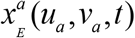 and 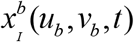 is given by the following neural network equations, known as the Wilson-Cowan Equations^34,35^

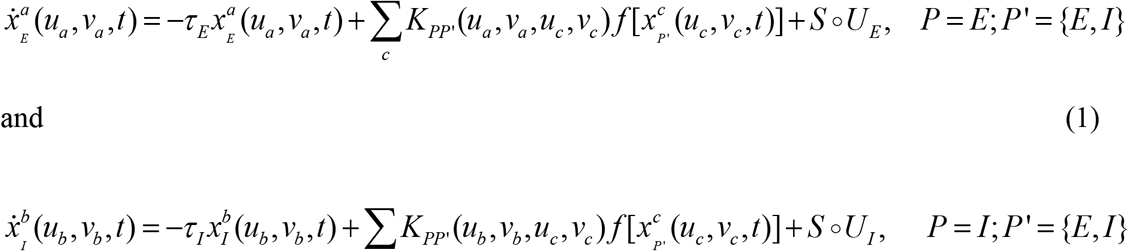

where *S*: ℝ → ℝ^*n*^ maps exogenous inputs to depolarization and *f* is vector-valued transfer function that describes the mapping from membrane potentials to current (spikes per second; Lipschitz continuous to guarantee local existence) of the population around point (*u_a_, v_a_*) ∈ *M_A_*.

We then take the continuum limit of Equations (1). This is a common transformation of biophysical evolution equations^36^ and allows one to replace sums with integrals. It follows a standard process in mathematical physics that provides the continuous version of a discrete system (opposite of discretization). We then partition *M_a_* into *N* × *L* cortical patches of neural densities 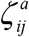 with dimensions (Δ*ν*, Δ*v*) *i* ∈1,…,*N* and *j*∈1,…,*L*. Thus the subgroup of neurons in the square 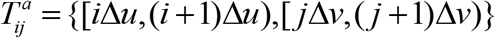 of *M_A_* is given by 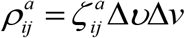. For mathematical convenience, consider a copy *M_B_* of manifold *M_A_*. The interaction between neurons in cortical patches 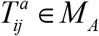 and 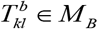 only depends on the duplets (*i, j*) and (*k, l*). A neuron at location (*u_a_, v_a_*) inside square 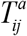 receives input from all neurons in square 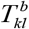 with strength 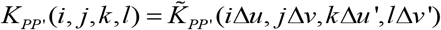, where we use “ ′ ” to denote locations on manifold *M_B_*. Also, 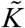 is the continuous version of function *K* under the assumption that connectivity is constant within the square with sides of length Δ*ν*and Δ*v*. For simplicity of notation, in the following we write *K* in place of 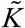. Then, we can define the local spatially averaged activity variable *X_P_* by 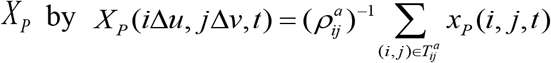 and consider the continuum limit Δ*u*, Δ*v*, Δ*u*’, Δ*v*’ → 0 : all nodes within the patches 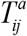 and 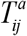 occupy the same location in manifolds *M_A_* and *M_B_*. After replacing *u* = *i*Δ*u, v* = *j*Δ*v* and *u*’ = *k*Δ*u*’, *v*’ = *l*Δ*v*’, Equations (1) can be written as a system

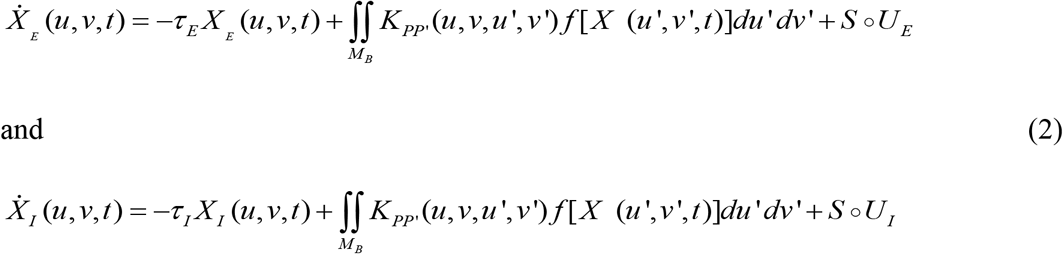

Similarly to ^29^, we then consider perturbations 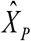 of membrane potentials around baseline: 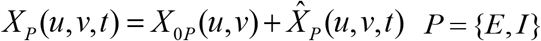. This yields an expression of the perturbations 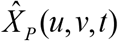 in terms of: 1) the functions *G_k_*, which we previously called *principal axes*^29^; and 2) the latent variables 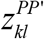, which we called *connectivity components*—to resemble standard PCA terminology. Both are defined below. In that earlier work^29^, we found that the principal axes contained temporal information, while the connectivity components contained spatial information. The connectivity components 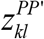 were defined by the following equations

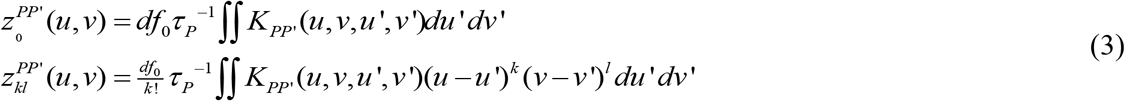

while the principal axes were given by

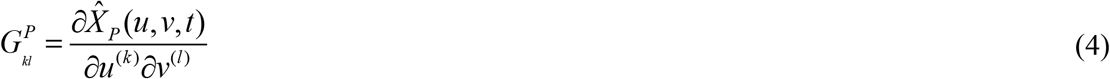

Using Equations (3) and (4), Equation (2) yields the following expressions for the perturbations 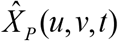:

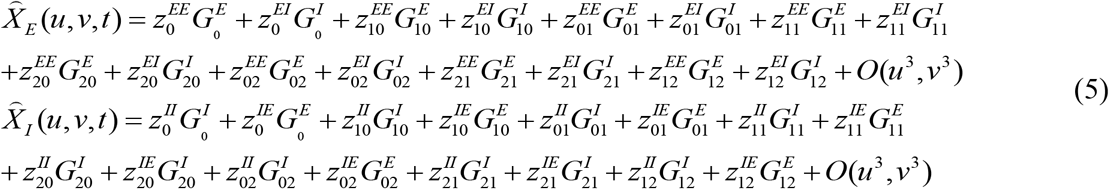

Note that the above equation is obtained using linear stability analysis and includes a Taylor expansion over spatial coordinates. If we had separate data (depolarization or spike rates) for the excitatory and inhibitory populations, we could use Equations (5) and this data to find 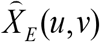 and 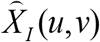 separately. We could estimate the connectivity components 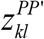 for the excitatory and inhibitory populations separately. We will pursue this in future work using data from excitatory and inhibitory neurons. Here, our data included aggregate activity (LFPs) from both populations.

LFP recordings contain aggregate activity of excitatory and inhibitory populations together. Mathematically, this is expressed as a two factor sum of membrane depolarization of all populations for each location on the cortical surface, 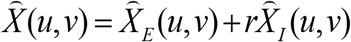, where *r* is the ratio of excitatory to inhibitory activity, which we take *r*=0.25. This value for *r* was chosen according to Dale’s principle that neurons can be either excitatory or inhibitory and there are four times more excitatory than inhibitory neurons ^37,38^. For mathematical convenience and without loss of generality we also consider a (differentiable) change of coordinates 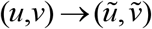 where 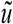 parameterizes the location of the excitatory populations and 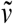 parameterizes the location of the inhibitory populations. We also assume that the Jacobian of this transformation 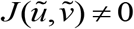. In the Results section, we validated this assumption numerically. The rigorous mathematical justification of this assumption will be considered elsewhere. Following this, the principal axes 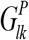 and components can be simplified:

i. 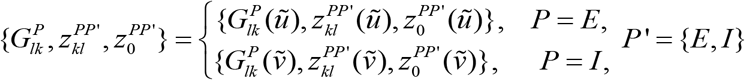
ii. 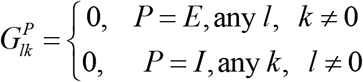

Thus: (i) Principal axes 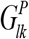 and components 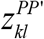 describing excitatory populations depend on 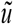 only and terms describing inhibitory populations depend on 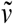 (this was the assumption above); (ii) Axes 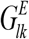 involving excitatory activity involving non zero sub-indices *k* can be removed from Equations (5), because these axes contain mixed derivatives. Similarly for inhibitory activity and its axes 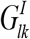 that contain mixed derivatives with non zero sub-indices *l*. Because 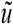 and 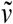 are distinct (the locations of excitatory and inhibitory populations are different), we can consider the union of the spatial domains for 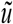 and 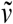 as a single, new spatial domain and join the spatial variables 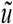 for the location of excitatory and 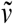 for the location of the inhibitory populations into a single variable. Then, adding Equations (5a) and (5b), we obtain

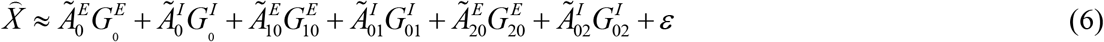

where the aggregate connectivity components 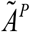 are two factor sums of *z^PP′^* defined by Equations (3):

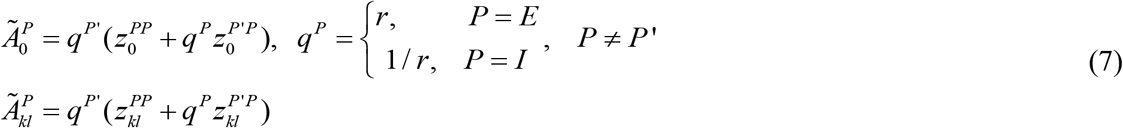

Letting 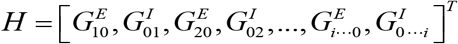. Equation (6) is a *deep neural field,* and can be rewritten in the general form of a Gaussian Linear Model (GLM; cf. Equation (1) in ^29^),

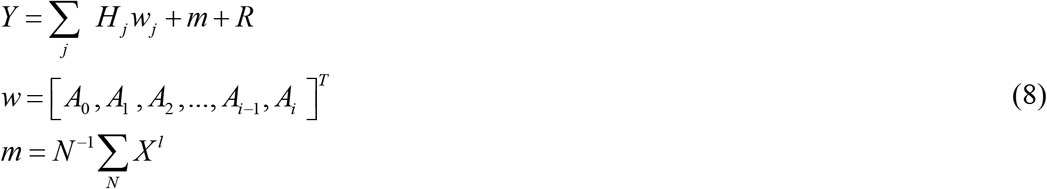

where for simplicity of notation we have relabelled, 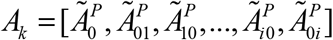 and have dropped the superscript *P*, because we do not distinguish between neural populations in what follows. This simply relabels components with two sub-indices as components with a single sub-index. Note that *A_k_* are 1D, while 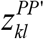 are 2D. Since there is only one spatial variable in *A_k_*, only one sub-index was needed. We have also assumed that cortical activity 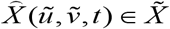 was sampled from a random process 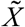 and 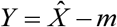.

The above 1D reduction was obtained under certain mathematical assumptions. To validate them, we compared our effective connectivity estimates against two established approaches (see Methods subsection below and Results section). We found that our results correlated significantly with results obtained with these methods. The rigorous mathematical justification of these assumptions will be pursued elsewhere.

To sum up, starting from a neural network model for coupled excitatory and inhibitory populations (Equation 1), we have shown how it can be reformulated as a deep neural field model (Equation 6) – and then a GLM (Equation 8). This is useful because it allows us to obtain the effective connectivity that characterises information flow within the neural ensemble. This is described in the next section.

### Connectivity components and kernels

The connectivity components *A_k_* are the latent states of the autoencoder trained by optimizing the cost function, known as the Free Energy, *F*,

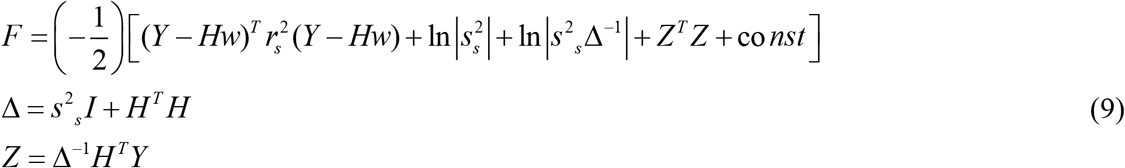

using a Restricted Maximum-Likelihood (ReML) algorithm^39^. This assumed a directed graphical model *p*(*Y*|*w*) used in autoencoders that yields an approximation *q* to the posterior *p* ~ *N*(*w*|*Y*), see ^29^ for more details. Note that the cost function defined by Equation (9), is the same cost function like the one used in Predictive Coding.

To sum up, Equations (3) define the connectivity components *z^PP′^* of the neural network (1). Similarly, Equations (7) define the 1D connectivity components *A_k_* of the deep neural field (Equation (6)) as two factor sums of *z^PP′^*. Training the GLM to optimize the cost function (9) we obtain single trial estimates of effective connectivity components *A_k_*. Their averages across trials are shown in Figure 2 of ^29^. If we had separate recordings of excitatory and inhibitory neurons we could get the effective connectivity components *z^PP′^* of the neural network (1) in a similar way. This will be pursued elsewhere. Here, we used single trial *A_k_* estimates to identify neural ensembles that maintained location during each trial. We also compared them to similar measures obtained using other approaches for ensemble identification (see Methods below and Results).

We now turn to connection weights of the neural network (1). We call these connectivity kernels *K_PP_*,. In Equations (3), the connectivity components are integrals of the connectivity kernel *K*(*u_a_, v_a_, u, v_b_, t, t*′). Here we have dropped the sub-indices *P, P’* because the kernel is not spatially discrete; instead, it depends on continuous variables (*u, v*).

In ^29^, after obtaining *A_k_*, we assumed that cortical connectivity has a Gaussian profile and computed 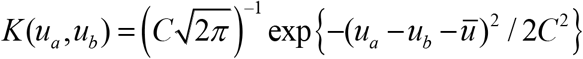 and obtained trial average estimates of *K*(*u_a_, u_b_*) where 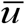 and *C* are the mean and standard deviation of axonal dispersion. Here, we first considered the same profile and focused on the corresponding single trial estimates of *K*(*u_a_, u_b_*). We considered a more general expression for the connectivity profile involving a weighted Gaussian (see *Mapping the latent space to a cortical patch* section below).

### Comparison of our approach to established approaches in the literature

To validate our approach, we compared our estimates of connectivity components and kernels to methods that are established in the literature. First, we considered a correlation-based method, see ^40^. This yields neuronal ensembles, where neurons in the same ensemble have dense connections with each other and weak connections to other neurons. This is achieved by maximizing a graph theoretic measure known as modularity and is similar to finding communities in social networks ^41^. It computes similarity measures including cosine similarity and the correlation coefficient that we used here. The method was initially developed to analyse spike train data, but we here adapted it to deal with LFPs. It provides a spectral decomposition of the modularity matrix using a stochastic algorithm ^41^. It employs a consensus algorithm to ensure that the same clustering is obtained for different initialisations ^42^. This method has been applied to neural activity in visual cat areas ^40^ and the Aplysia pedal ganglion ^43^.

Second, we used a higher dimensional SVD method known as canonical decomposition (CD^44,45^). This provides a generalization of the usual SVD which factorizes a tensor in terms of *R* arrays. It allows one to obtain an approximation of the data represented by a third order tensor *Y* ∈ ℝ^*N_r_*×*N_S_*×*T*^ given by

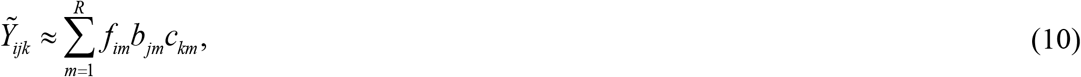

where *f_im_* ∈ ℝ^*N_T_*×*m*^, *b_jm_* ∈ ℝ^*N_S_*×*m*^ and *c_km_* ∈ ℝ^*T*×*m*^ are three matrices known as “modes” in the mathematical literature ^45^. Their first dimensions are either number of trials (*N_T_*), or electrodes (*N_S_*) or time (*T*). *R* is known as the rank of *Y*, with *m*=1, …*R*. Equation (10) can be written explcitly as 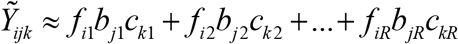. This includes a sum of combinations of elements *f*_*i*1_, *b*_*j*1_, *c*_*k*1_, *f*_*i*2_, *b*_*j*2_, *c*_*k*2_ Taking together (i.e. for all *m*=1,…*R*) all elements with the *same* first dimension, e.g. the dimension denoted by index “*i*”, that is,*f*_*i*1_,.*f*_*i*2_,.., *f_iR_* we obtain a matrix *f* = [*f_im_*] and similarly for *B* = [*b_jm_*] and *C* = [*c_km_*]. *F, B* and *C* are known as *modes* Each mode is a matrix where the first dimension (denoted by *i, j* or *k*) is equal to one of the above three dimensions of the LFP array, that is, a number of trials (*i*=1,..,*N_T_*), electrodes (*j*=1,…,*n_s_*) or time points (*k*=1,…, *T*). Thus, each term in the sum 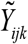 is a product of elements from the three modes *f_im_ b_jm_ c_km_*. This product is called a *factor.* The second dimension (denoted by *m*) is the same for all three modes that belong to the same factor and is different for each term in the sum (i.e. each factor). It ranges between 1 and some arbitrary number *R, m* = 1,…,*R*. Thus, *R* is equal to the number of factors in the CD approximation 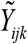. In the Results section, we will see that *R* can be estimated based on some measures from statistics.

The approximation 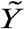 is obtained using an alternating least squares algorithm (ALS) that minimizes the reconstruction error 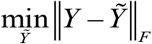, where ||*Y*||_*F*_ is the Frobenius norm of *Y*. The ALS approach fixes *B* = [*b_jm_*] and *C* = [*c_km_*] to find *F* = [*f_m_*]. The conditional least square estimate of *A* is then

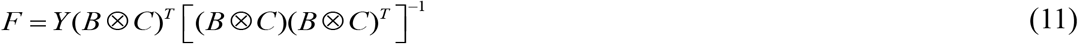

where ⊗ is known as the Kronecker product. ALS continues by then fixing *F* and *C* to find *B* and finally *F* and *C* to find *B.* The CD approximation is unique up to permutation and scaling of the modes. Thus, ALS is used iteratively. For more details see ^46^. CD has recently been applied to analyse spiking data and identify neural ensembles in ^47^. Here, we adapted this work to identify neural ensembles using LFPs. The CD approach by ^47^ does *not* provide single trial estimates of connectivity components and kernels, like those we considered here. However, we compared our results to CD estimates after averaging across all trials that corresponded to the same stimulus.

In the expansion (10) above, the number of components is arbitrary. To find the rank *R,* we used two criteria: consistency and congruence. Consistency was introduced as an alternative way to obtain the rank in CD approximations ^48^. It uses certain elements of CD theory, known as CD factors, to compute an alternative approximation of the data matrix, known as Tucker3 approximation ^49^. The Tucker3 approximation also contains (mixtures of) CD factors. For a given *R,* consistency quantifies the difference between data fits using the CD and Tucker3 approximations. *R* should be such that this difference is minimal. According to ^48^, this corresponds to consistency values between 50-100%. To sum up, consistency quantifies the degree that the LFP data contain a trilinear variation, see ^48^ for more details. It is optimal for that particular value of *R,* that renders the core of the corresponding Tucker3 approximation (the Tucker3 approximation with the same CD factors) superdiagonal.

Congruence, on the other hand, is simply based on uncorrected correlation coefficients (CC) between any two sets of factor matrices {*F_1_, B_1_, C_1_*} and {*F_2_, B_2_, C_2_*}. These are averaged over a different implementations of the ALS algorithm starting from different initial conditions and then the maximum value is subtracted from 1, i.e. congruence (CG) is given by 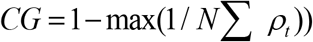 where *ρ_t_* is the CC computed in the *t*-th initialisation and we have assumed *N* initialisations. Congruence was initially used to remedy instabilities and slow convergence that are knowns to affect ALS, due to its iterative nature. A low value of congruence implies that the CD approximation was *not* stuck in local minima and CD factors are stable ^50^. In Results, we chose a rank *R* with high consistency and low congruence. This results in a stable CD approximation that includes a trilinear variation in the data.

### The electric potential and electric field generated by a neural ensemble

To model the *electric potential* (EP) generated by synaptic activity (EPSPs and IPSPs) in a neural ensemble we use the bidomain model of the neural tissue ^31^. This assumes that the neural tissue can be represented by a cylindrical fiber of radius *d*. This means that the problem has rotational symmetry and the potential is a function of two coordinates (*ρ, Z*), where *z* = *u_a_* is the coordinate along the fiber axis and *ρ* is coordinate vertical to it, see ^51^ and Supplementary Figure 4. Below, we use the bidomain model, to derive the extracellular potential *V^e^* and the extracellular electric field generated by the neural ensemble, *E^e^*. Details of this derivation can be found in the references above. Here, we included a summary for the convenience of the reader. The bidomain model describes the potential in the two sides of the neuron membrane, that is, the intracellular *V^t^* and extracellular *V^e^* potentials. Their difference 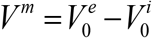 is the transmembrane potential and results in a spatial discontinuity also for the electric field 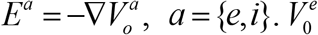 and 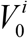 are the values of the extracellular and intracellular EPs on the two sides of the membrane. Note that ∇ denotes the gradient operator. According to the theory of electromagnetism, this discontinuity gives rise to dipole sources with moments ^27^

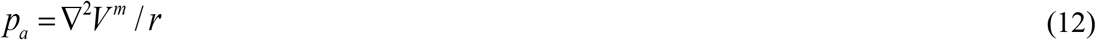

Here *r* is the brain resistivity with *r* = 2.2 Ohm^52^ and we have assumed that the number of neurons is large and that each cell is very small compared to the distance at which the LFP electrode is placed. Also, the current density *I^a^* (*u_a_, v_a_*) that results from EPSPs and IPSPs is given by

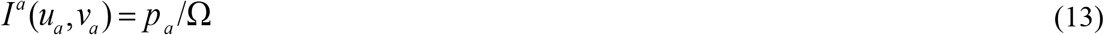

where Ω is the total volume of the ensemble. Neglecting ephaptic interactions *V^m^* ≈ *V^i^*, and the extracellular electric potential generated by the current density *I^a^*(*u_a_, v_a_*) is given by

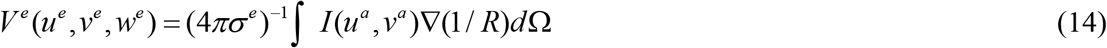

where *σ^e^* is the conductivity of the extracellular space, and *R* is the distance between the current source at the point (*d, u^a^*) of the neural ensemble and the point (*ρ, u^e^*) in the extracellular space where we measure *V^e^*, i.e. the location of the LFP electrode, 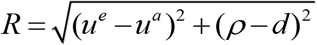, see Supplementary Figure 4. Then, according to the bidomain model, Equation (14) can be written as ^53,54^

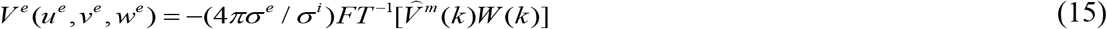

where 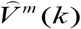 is the Fourier Transform of the transmembrane potential *V^m^* and *FT^-1^* is its inverse Fourier Transform, that is,

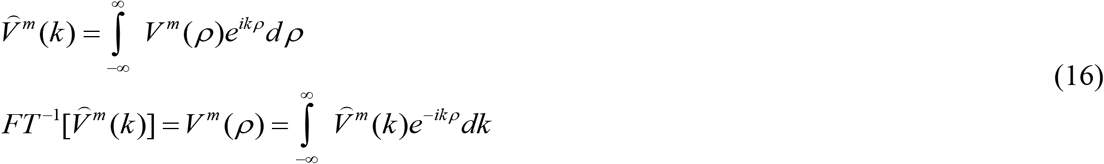

The function *W*(*k*) is given in terms of the modified Bessel functions of the first *I*_0_(*ρ*), *I*_1_(*ρ*) and second *K*_0_(*ρ*), *K*_1_(*ρ*) kind ^55^,

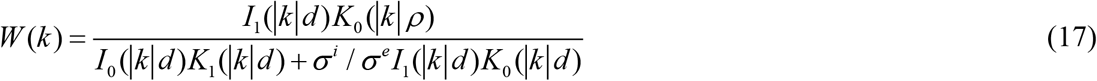

Then, the *extracellular electric field* (EF) generated by the neural ensemble, *E^e^*, is just the gradient of *V^e^*, *E^e^*. –∇*V^e^*

### Gauge transformations of electric potentials

Multiple extracellular EPs *V^e^* can give rise to the same EF *E^e^* = –∇*V^e^* in extracellular space. This is a well-known result in the theory of electromagnetism called *Gauge invariance.* It follows from the conservation of electrical charges ^27^. In the case of LFP measured with multielectrode arrays, each trial gives rise to an LFP recording. This, in turn, results from a different EP generated by current flow within a neural ensemble in each trial. In Results, we test the *hypothesis* that the *EF is the same for all trials corresponding to the same remembered stimulus, E^e^*{ trial *i*} = *E^e^* { trial *j*}. To test this hypothesis, we first needed an estimate of the extracellular potential (EP) at an arbitrary trial *j, V^e^*{ trial *j*}. We obtained this using Equations (15) and (17) above, in two ways. First, using simulations of our deep neural field model. Second, using recorded LFPs as proxies for the transmembrane potential at arbitrary trial *j, V^m^*{ trial *j*}. Then by taking the gradient of *V^e^* { trial *j*}, we found the extracellular EF for trial *j, E^e^*{ trial*j*}. Having obtained EF estimates, we tested the hypothesis that the EF is stable in three ways: First, we looked whether EFs where correlated across trials. Second, we asked if EF estimates were consistently different for neural ensembles that maintain different cued angles. We tested if we could distinguish between memorized cues based on EFs. We used EFs as classification features in two commonly used classification algorithms, Naïve Bayes and diagonal LDA^29^. Third, we used Gauge functions that connect the recorded LFPs. If the EF was stable, the EPs are related by called *Gauge transformations* ^27^

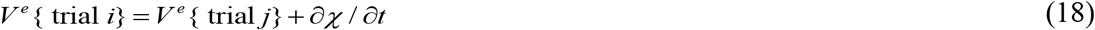

where the function *χ* = *χ*(*ρ, ζ, t*) is called *Gauge function.* To sum up, a third way to test the stability of EFs is to test if the Gauge functions can be used to distinguish between different cued angles (see Results). According to Equation (18), the time derivative of the Gauge function *∂χ*/*∂t* is equal to the difference of EPs corresponding to any two trials. Equation (18) should hold for any arbitrary pair of trials. Thus, we asked whether we could decode cued angles using Gauge function derivatives *∂χ*/*∂t* as classification features. These, in turn, were obtained after subtracting LFP recordings. An independent experimental validation could also be carried out using intracellular recordings: If Equation (18) holds, then a similar Equation for the intracellular potential *V^t^* also holds with the same Gauge function *χ* = *χ*(*ρ,ζ, t*). Thus, the Gauge function *χ*(*ρ, ζ, t*), can be found experimentally by measuring *V^i^* during any two trials, *V^i^* { trial *i*} and *V^i^* { trial *j*}.

### Mapping the latent space to a cortical patch

The extra step that allowed us to obtain the electric field above was the mapping of the latent space to a cortical patch ^29^. Starting from the connectivity components, we obtained the weights that scaled incoming input to each population from all other populations in the ensemble, called the connectivity kernel. This describes information exchange and electrical activity on the patch. Having this, we then reconstructed the EF. Consider Equation (1). The connectivity kernels *K_PP_*,(*u_x_, v_x_, u_e_, v_e_*), *X* = {*a, b*} include the weights that scale input from a population at location (*u_c_, v_c_*) to an excitatory population at (*u_a_, v_a_*) or an inhibitory population at (*u_b_, v_b_*). Above, we considered the continuum limit of Equations (1), that is, Equations (2) and similarly the continuum limit of the connectivity kernels*K_PP_*,(*u, v, u*′, *v*′). These have the same meaning as *K_PP_*,(*u_x_, v_x_, u_e_, v_c_*). Only a difference in notation: the sub-indices denoting location have been replaced by continuous spatial variables that lie on a patch *u, v* ∈ *M_A_*. Then, given the connectivity components *A*_0_, *A_kl_*, we can find *K_PP_*,. In mathematical terms, the kernels are probability distribution functions and can be estimated using a variety of methods from inverse problems theory, including splines ^56^, series expansions ^57^ and other methods ^58,59^.

We here considered a Gaussian connectivity profile used in ^29^ and an alternative expression for the connectivity kernel that includes sums of Gaussian profiles weighted by polynomial factors known as Hermite polynomials, *H_n_* ^55^. These sums are known as Gram-Charlier series. In brief, the connectivity kernel of the neural ensemble can be approximated by

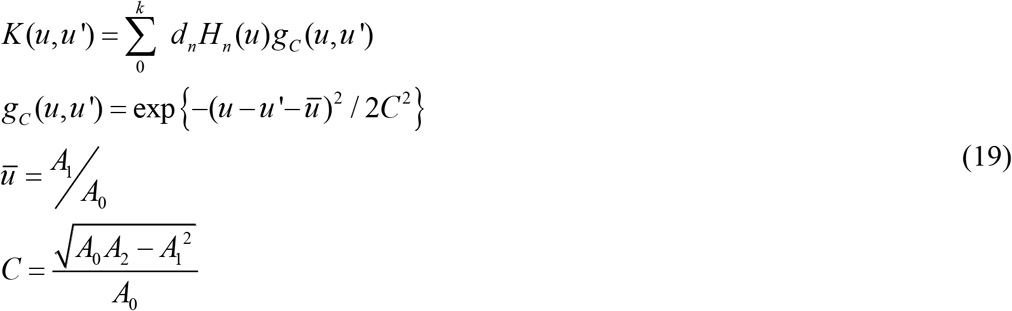

where the Hermite polynomials, *H_n_* are *known* and the coefficients *d_n_* can be found by substituting (19) and the definition of *H_n_* into

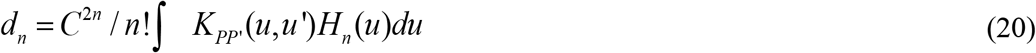

Interestingly, Equation (20) using the binomial theorem and the definition of connectivity components gives

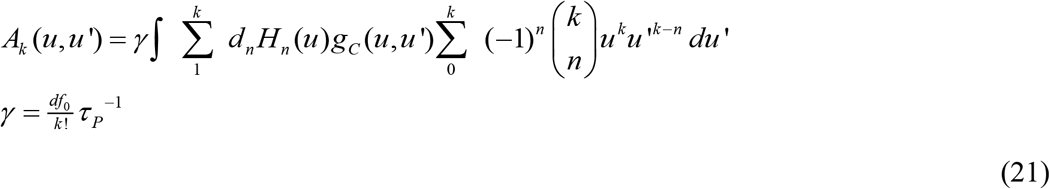

The above expression seems complicated. However, one can use the properties of the Hermite polynomials to find the coefficients *d_n_, n* = 0,1,…,*k*.

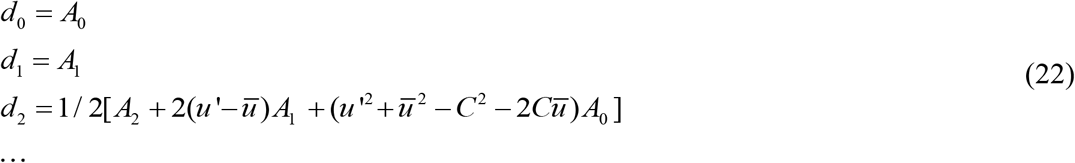

Substituting the above expressions and the expressions for Hermite polynomials into Equation (19), we obtain an alternative expression for the connectivity kernel *K*(*u, u*′) that involves a Gaussian function weighted by terms involving connectivity components (keeping the first three terms in the series expansion given by Equation (19)):

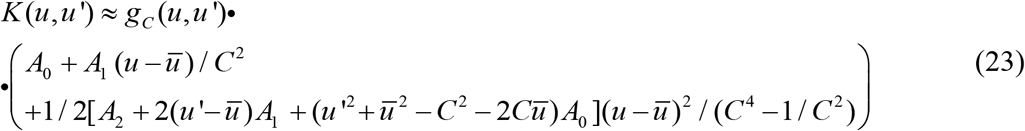

## Results

### Deep neural fields describe neural ensemble structure in a holistic fashion

This paper follows upon our recent work that focused on groups of neurons that represent memories known as neural ensembles (Figure 1). In ^60^ we studied computations performed by neural ensembles during a flexible sensorimotor decision making task^61^. We showed that neural ensembles in the same brain area performed different computations based on the rule applied during each trial, although the stimulus processed was the same. This result was obtained by comparing brain responses to both a behavioral model and a deep neural network and testing if they give similar results. In a parallel line of work^29^, we also studied the structure of neural ensembles and obtained their effective connectivity. Our analyses below build upon that earlier work and used the same dataset. This includes a spatial working memory task, where the angle of a cue had to be remembered (delayed saccade task; Supplementary Figure 1A). We analysed LFP data recorded during the delay period.

**Figure 1.**
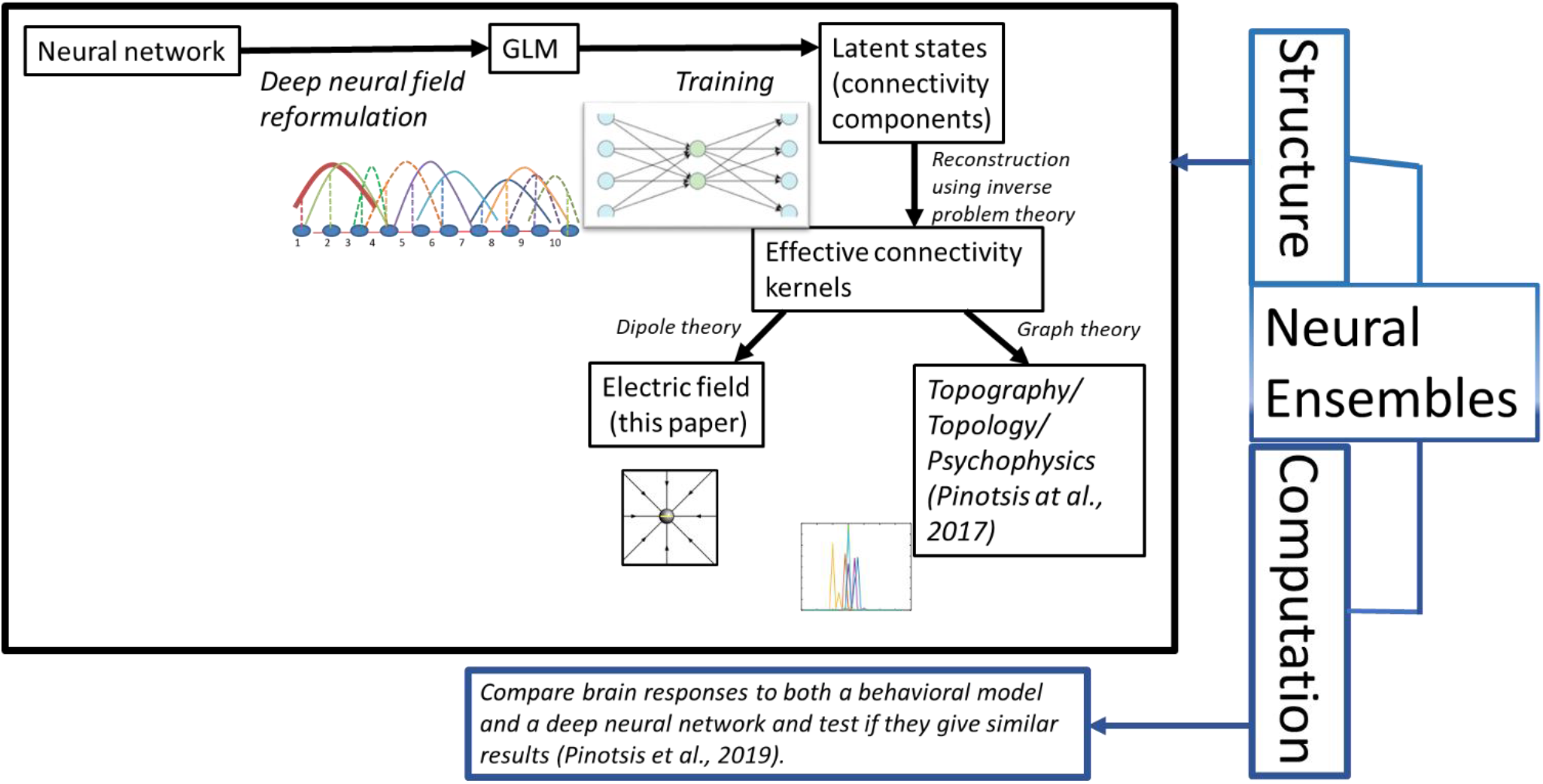
**A.** Outline of our approach. We first reformulated a neural network (described by Wilson Cowan equations) as a neural field model and then a Gaussian Linear Model (GLM). We trained this model as an autoencoder and obtained the latent states (connectivity components). Then, using inverse problem theory, we obtained the corresponding connectivity kernels. In ^29^, we used the kernels and graph theory to characterize the topography and topology of neural ensembles. Here, we use the kernels and electromagnetism (dipole theory) to study the stability of the electric field generated by an ensemble. This paper and ^29^ focus on the structure and biophysics of neural ensembles. In related work ^60^, we also studied the computations performed by ensembles using deep neural networks and behavioural models.

We analysed neural activity (LFPs) recorded from a multielectrode array of *N_S_* = 32 electrodes implanted in the FEF of two macaque monkeys. LFPs are thought to describe neural activity from a population in the proximity of each electrode ^62,63^. Analysing LFPs allowed us to identify neural ensembles and test if they overlap in different trials. Electrodes were numbered in a monotonic fashion; neighbouring electrodes had adjacent numbers (Supplementary Figure 1B). Our approach and the main results of ^29^ are summarized below and in Figure 1.

Our approach has the following steps: 1) Start with a neural network model. 2) Reformulate this as a biophysical Gaussian Linear Model (GLM), that we called deep neural field. The term “deep” was used in our earlier work^29^ to distinguish this model (with learned connectivity parameters) from common neural field models where connectivity weights are chosen ad hoc, e.g.^64–67^. The learned parameters are obtained after training the neural field as an autoencoder (see also ^29^). Thus, the term “deep” refers to the hidden layer of the corresponding training network. 3) Use the latent states (connectivity components) and inverse problem theory to obtain the effective connectivity (connectivity kernels). We will come back to components and kernels (and their differences) below.

In^29^, we used average connectivity estimates and graph theory and showed that path length portioned the space of cued angles. The smallest values occurred for cues on the horizontal meridian, i.e. information propagates faster. This provided an explanation of the oblique effect in psychophysics^68^. Here, we use single trial connectivity estimates and dipole theory to reconstruct the electric field produced by a neural ensemble. This will be discussed in detail later.

Examples of neural activity for two different individual trials corresponding to the same task condition are shown in Figure 2A. LFP amplitudes (in *mV*) are shown on the vertical axis. The horizontal axis are electrode number (location) and time (in *ms*). We assumed that FEF comprised a large number of neural populations (indexed by *j*=1,.,.,*N*=32), that was equal to the number of electrodes we sampled from, see also ^29^. Each of these populations can be thought of as centred around a point (*u_a_, v_a_*) on the 2D cortical surface. They also interact with other populations located at point (*u_b_, v_b_*), via an effective connectivity kernel *K* (*u_a_, v_a_*, *u*_b_, *v*_b_, *t, t*′).

**Figure 2.**
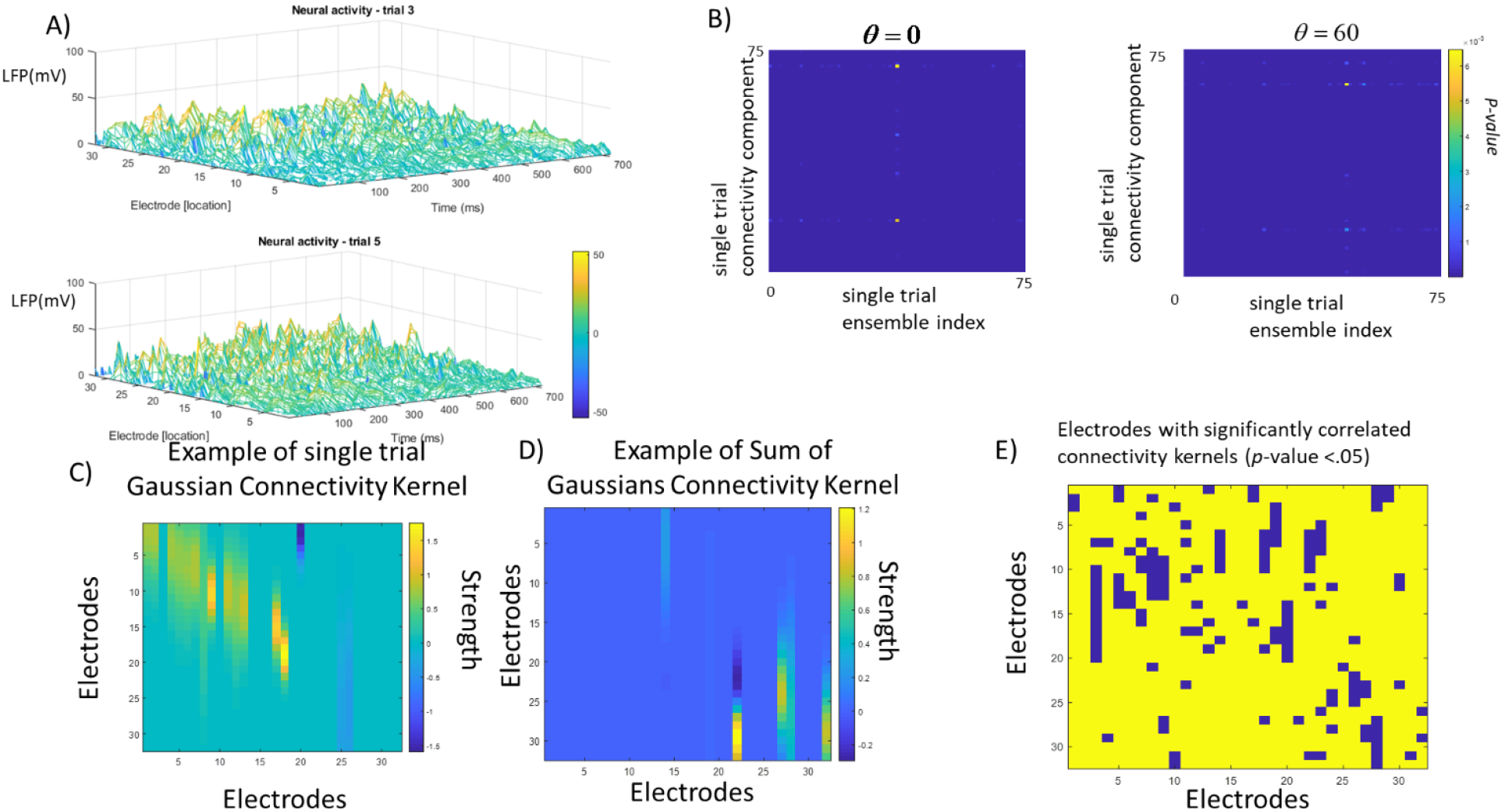
**A.** Examples of neural activity for two different individual trials corresponding to the same task condition. Local field potentials (LFPs, in *mV*) are shown on the vertical axis. The electrodes (location on the cortex) and time (in *ms*) are shown on the two horizontal axes. **B.** Significance (*p*-value) of Pearson correlations between the single trial connectivity components and ensemble indices obtained by the approach of ^40^ Trials where a horizontal location was maintained (*θ*=0 degrees) are shown in the left panel. Similarly, trials for cued angle at *θ*=60 degrees are shown in the right panel. Estimates for all trials correlated perfectly (*p*<10^-2^). **C.** Example of effective connectivity kernel with a Gaussian profile. This describes the weights which scale neural activity propagating between any pair of populations located near one of the electrodes. This kernel characterizes information flow at the single trial level. **D.** Example of an alternative expression of the effective connectivity kernel obtained as a weighted Gaussian using a series expansion. **E.** Correlations between the connectivity kernels in panels C. and D. *R*=87% of connectivity weights were significantly correlated at the *p*<.05 level. These are shown in yellow. Blue denotes weights that were not significantly correlated.

We previously identified neural ensembles based on their effective connectivity kernel averaged across trials ^29^. This connectivity was expressed in terms of two measures: 1) the latent variables of an autoencoder that we called *connectivity components* and 2) the *connectivity kernel K* (*u_a_, v_a_*, *u*_b_, *v*_b_, *t, t*′) of a biophysical rate model (neural field). This kernel was obtained from the connectivity components after assuming a Gaussian connectivity profile over space. Here, we followed a similar approach and focused on effective connectivity of a neural ensemble and its components at the single trial level (i.e., without averaging). We also considered a more general weighted Gaussian as a connectivity profile over space. Our starting point was different to ^29^: we modelled each neural ensemble as a 2D neural network model of interacting excitatory and inhibitory populations (Wilson-Cowan Equations; see Methods). By changing the variable that parameterised the cortical surface from discrete to continuous, the neural network was reformulated as a mean field model, known as a neural field ^34,69,70^. In ^29^, our starting point was a usual neural field.

Since we are measuring aggregate activity (LFPs), we could not distinguish between locations of excitatory vs inhibitory populations. At the same time, these locations do not overlap. Intuitively, this means that we can join the 2 spatial variables in the neural network, describing locations of excitatory and inhibitory populations, into one. To sum up, the original 2D neural network model was first transformed to a 2D neural field and then to an 1D deep neural field considered in ^29^. The details of this reduction are included in Methods. Its mathematical implications will be considered elsewhere. Here, we assessed whether this reduction allowed us to correctly identify neural ensembles, by comparing our results to those obtained using other methods.

### Effective connectivity components of deep neural fields reveal clusters obtained using pairwise correlations

We compared the effective connectivity components and kernels obtained with the deep neural field model to estimates of ensemble connectivity obtained with other methods. Below, we show that our effective connectivity estimates correlated significantly with connectivity estimates obtained using pairwise correlations ^40^ and a high dimensional SVD approach ^44^.

We first discuss connectivity components (denoted by *A_k_* in Methods). These are the latent variables of the low dimensional space obtained after training our deep neural field as an autoencoder. We will see below that they describe aggregate synaptic input to neural populations located at a certain point on the cortical patch. They cluster neurons into task related groups ^47^. Specifically, we obtained single trial component estimates in the following way. We trained a deep neural field model using a cost function considered in predictive coding and autoencoder networks. Component averages across trials were shown in Figure 2 of ^29^. In that paper, we also showed that connectivity components were matrix-valued functions with dimensionality equal to *N*_T_ × *N_S_*, where *N_T_* = 600 is the number of trials. For each trial, we obtained a vector of dimension *N_S_* whose entries were called *component strengths*. These were similar to loadings or principal components in PCA. Because we here used aggregate neural activity from both kinds of populations (LFPs), we obtained effective connectivity components for both populations together. This is similar to other dimensionality reduction approaches, e.g. ^8,47^.

Here, we validated the effective connectivity components obtained in ^29^ (summarized also in ^71^) using two independent methods. First, using a correlation-based method, see ^40^. This was originally used to identify similarities between spike trains. It was based on pairwise correlations. Similarities were then used to define neural ensembles –assuming that neurons with similar spiking patterns represented the same stimulus or sequence. Thus, one obtains neural ensembles. Each neuron is included in an ensemble (called a “cluster” in the original paper), indicated by an ensemble index. In other words, the approach by ^40^ did not yield effective connectivity per se, but one can map the ensemble index to effective connectivity components and kernels that we obtained.

Below, we compare our effective connectivity estimates to the clusters obtained using the method presented in ^40^. This yields an alternative way to obtain the same neural ensembles described by our deep neural field model in an unsupervised way, using a *k*-means algorithm. We adapted the original algorithm from ^40^ to work with LFP data, instead of spike trains. For each trial, the method assigned each electrode to an ensemble using an ensemble index. Assuming that electrodes sample from populations in their proximity ^62,63^, this process also assigns populations to ensembles. We then computed the correlation between the ensemble indices and the first connectivity components for different cued angles (angles). We asked whether the ensemble index correlated with the component strength for each electrode and trial. In ^29^, we showed that the component strengths are aggregate sums of all the weights of all connections that target the electrode at hand. They describe changes of signal as it propagates between electrodes. Thus, different values of component strengths correspond to different levels of activity (drive) that each electrode receives.

Pearson correlations were obtained for ensembles obtained from trials with different cued angles. Recall that the monkey performed a spatial delayed saccade task (Supplementary Figure 1A). The *p*-values across all 32 electrodes are shown in Figure 2B for a remembered stimulus at angle *θ* = 0 (left) and *θ* = 60 (right) degrees. Correlations were also significant for trials that involved different angles (other stimuli, not shown). It should be noted that both the ensemble index and the component strength of the deep neural fields are single trial measures. The fact that they were significantly correlated implies that electrodes formed ensembles based on the drive that the neural population in the vicinity of each electrode received during each trial. This is similar to intercolumnar synchronization observed in perceptual grouping studies^72^.

To sum up, we found that the effective connectivity components of our deep neural field model describe the same clusters as those found using pairwise correlations ^40^. |This provides an independent validation of our effective connectivity estimates at the single trial level.

### Connectivity kernels correlate with ensemble indices

Recall that, besides connectivity components, our approach also yields the connectivity kernel. Its entries, called *connectivity weights* quantify the strength of the effective connections between the recording sites within each cortical area, see Supplementary Figure 1B. They multiply input signal from other electrodes that targets a certain electrode measuring activity from a part of the neural ensemble. In other words, they describe how the signal is amplified or attenuated when it propagates between recording sites. Large positive weights of connections targeting a certain electrode implies that large LFP responses would be expected from that recording site.

The connectivity kernel enabled us to map the latent space (spanned by the components) to a cortical patch, that the neural ensemble occupies. Later, we will also use the connectivity kernel to predict the electric field generated by the ensemble. First, we assessed whether the connectivity kernel could also identify neural ensembles, similarly to the connectivity components above.

We assumed a Gaussian connectivity profile over space and obtained single trial connectivity kernel estimates *g_c_*(*u_a_, u_b_*) (Methods). We also considered a more general weighted Gaussian profile. This is a generalization of the widely used Gaussian kernel^64,65,67^ that follows from a series expansion ^57^. Here, a Gaussian kernel is weighted by known Hermite polynomials ^55^.

Example of these connectivity kernels obtained using data from a random trial are shown in Figure 2C and 2D. Figure 2C shows a single Gaussian kernel, while Figure 2D shows the more general weighted Gaussian. Note that because of the Gaussian profile, only elements around the main diagonal are non-zero. Figure 2E shows correlations between the two expressions obtained. *R*=87% of connectivity weights of the kernels shown in Figures 2C and 2D were correlated.

For simplicity, in the analyses below we used the expression involving a single Gaussian kernel. Similar analyses can be carried out using alternative expressions. First, we asked whether the connectivity kernel could be used to identify neural ensembles, similar to the analyses for connectivity components presented above. We computed correlations between single trial connectivity kernels and ensemble indices, obtained using the method of ^40^.

Earlier, we found that ensemble indices were correlated with connectivity components.. Correlations were significant for all trials. This implied that electrodes formed ensembles, where electrodes in the same ensemble had neural populations in their vicinity driven by the same input. Similarly, we found that ensemble indices also correlated with the connectivity *kernels* we obtained. Correlations between ensemble indices and connectivity kernels for cued angle at *θ* = 240 degrees are shown in Figure 3A. Correlation coefficients are shown in the vertical axis and trials in the horizontal axis. Trials with significant correlations at *p*<0.05 are denoted with asterisk. Overall, 25-40% of single trial kernel estimates correlated with ensemble indices for different angles. The percentage of significantly correlated trials for each cued angle is shown in Figure 3B. Connectivity components were correlated with ensemble indices across all trials. Connectivity kernels across some of the trials where the same cued angle was maintained, not all. Note that the connectivity kernels were a priori constrained to have a Gaussian (parametric) form, while the components were unconstrained. This explains why the percentage of significant correlations is smaller in the case of kernels. Some ensemble indices show Gaussianity too – but there is nothing intrinsic in the method of ^40^ that requires this assumption—which, on the other hand, was intrinsic to the Gaussian profile we assumed for kernels.. If ensemble indices are not Gaussian, there are no significant correlations.

**Figure 3.**
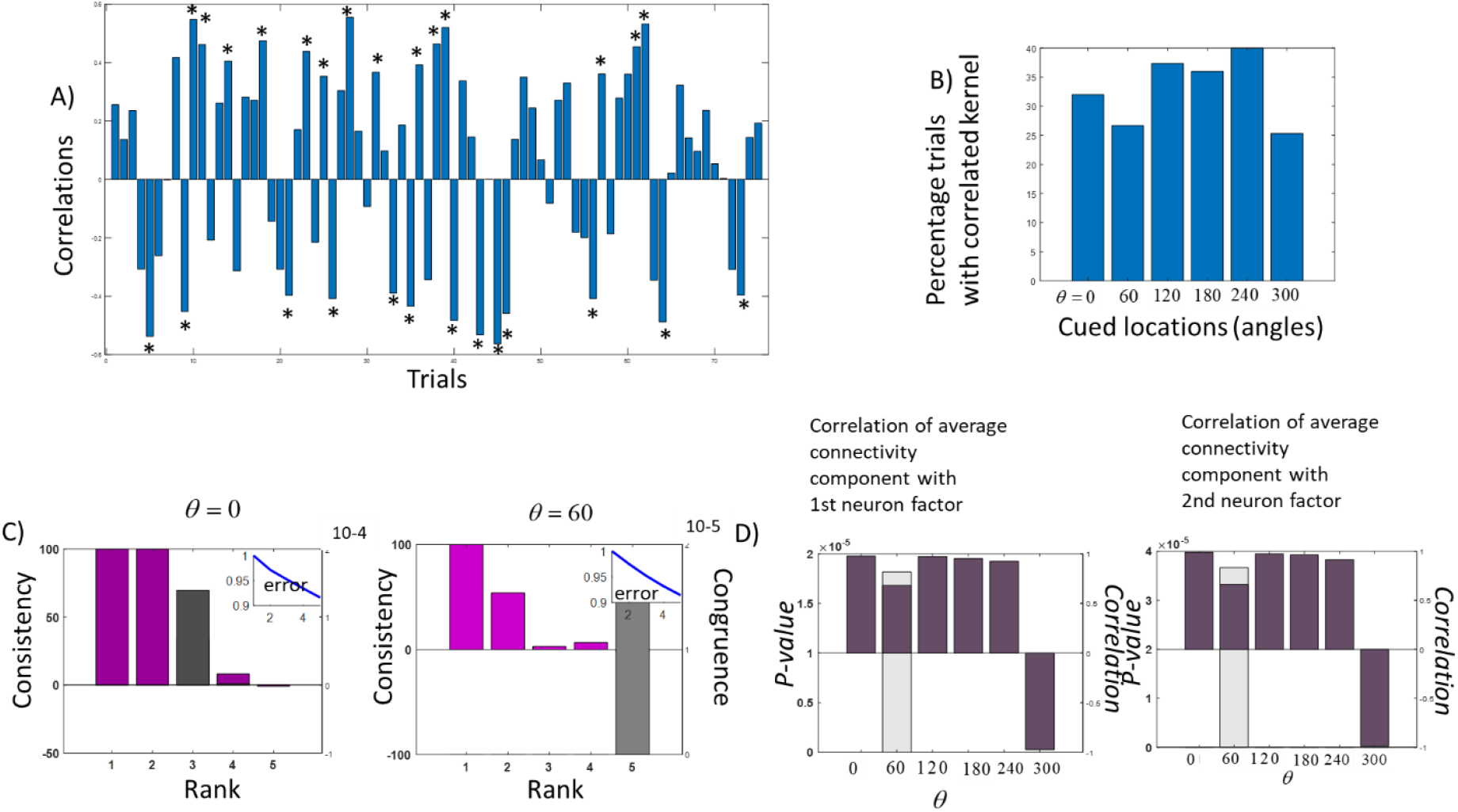
**A.** Correlations between single trial ensemble indices and connectivity kernels for cued angle at *θ* = 240 degrees. Correlation values are shown on the vertical axis. Individual trials are shown on the horizontal axes. Trials with significant correlations at *p*<0.05 are denoted with an asterisk. **B.** Percentage of significantly correlated trials for each cued angle. Locations are shown on the horizontal axis. Overall, 25-40% of single trial kernel estimates correlated with ensemble indices for different angles. **C.** Canonical Decomposition. Left and right panels show results for cued angles at *θ*=0 and *θ*=60 degrees. The number of factors (rank) is shown on the horizontal axis. Consistency is shown using magenta bars, while congruence is shown using grey bars. Different bars correspond to different ranks. Consistency values are shown on the left vertical axes, while congruence values are shown on the right vertical axes. ALS algorithm reconstruction error is shown in the insets. **D.** Correlations between connectivity components and first (left panel) and second (right panel) neuron factors obtained via Canonical Decomposition. Cued angles are shown on the horizontal axes. *P*-values (grey bars) are shown on the left vertical axes. Correlation coefficients are shown on the right vertical axes (burgundy bars).

All in all, the above results suggest that the connectivity kernels identified the clusters obtained with the pairwise correlation method of ^40^. The advantage that these kernels have over the previously considered components is that they describe actual connectivity on a cortical patch, not latent space.

### Connectivity components of deep neural fields correlate with high dimensional SVD components

We also validated the effective connectivity components obtained using our deep neural field approach using a second method. This is based on some old extension of high dimensional SVD, known as Canonical Decomposition (CD), see ^44^ and ^47^ for a recent application. Recall that, to obtain effective connectivity estimates, we trained the biophysical model as an autoencoder. This is similar to classical principal component analysis (PCA): Obtaining the connectivity components amounts to obtaining principal components. Thus, another validation of our components can be achieved by comparing them to components obtained using an SVD approach like CD. Note that CD components do not correspond to single trial ensemble connectivity like the components obtained using our method. CD provides an estimate of average (across trials) connectivity that we had found in ^29^. The authors of ^47^ called this average the neuron mode and suggested that it describes the “*spatial structure that is common across all trials*”. Below, we will see that this is similar to the average connectivity component across trials. We will also compare the CD neuron mode with the average connectivity component. We will show that the two correlated perfectly.

CD yields an approximation of our data: this is a three-dimensional LFP array *Y* ∈ ℝ^*N_T_*×*N_S_*×*T*^ (trials x electrodes x time; Methods). The CD approximation includes three matrices (or “modes”) that describe patterns over each of the three dimensions: a trial, electrode and time mode. These are dominant patterns in the data, similar to PCA components that describe dominant patterns in time or space. Examples of such PCA components include those obtained with the dimensionality reduction approach of ^73^ that outputs motor timing, i.e. trajectories in a low dimensional domain spanned by PCA components in the time domain; also in ^74^ PCA components included trajectories traced out by neurons representing motion and color.

We used the CD approximation to validate our effective connectivity estimates. Of particular interest for our current analysis is the neuron mode. This is an *N_S_* × *R* matrix, where *N_S_* is the number of electrodes and *R* is a constant, known as rank, that will be estimated below based on some measures from statistics. We assume that each electrode measures activity from a neural population in its proximity. The columns of this matrix are vectors of dimension equal to the number of electrodes. Each entry is an approximate LFP measured at each electrode (averaged across time and trials). The paper ^47^ used spiking data and the CD approach to obtain the neuron mode. These authors suggested that one can think of the neuron mode as a prototypical firing rate across neurons.

According to ^47^, a neuron mode corresponds to “*the synaptic weights from each latent input to each neuron”.* This is the *same definition* as that of the component strengths included in ^29^: *“(component strengths) express the sum of all connectivity weights that target the neurons that contribute to the LFPs observed from each electrode*”. Thus, our connectivity components and CD neuron modes are generalisations of principal components in three dimensions, and they have similar definitions.

Note that here, we used the word “mode” instead of “factor”, because the word “factor” is commonly used in the mathematical literature to denote terms in the CD approximation ^45^. In other words, our “neuron mode” is the “neuron factor” of ^47^.

We asked whether our connectivity components and CD neuron modes that can be found using our LFP data were correlated. We considered connectivity components averaged across trials. To find the CD neuron modes we used a standard iterative Alternating Least Squares (ALS) algorithm ^46^. Before obtaining the modes we needed to find the rank, *R* (Methods). This is part of the CD approach. We assumed different values for rank *R*=1,…,5. For each value of *R*, we calculated the sum of squares reconstruction error. This is plotted on the vertical axis appearing in the top right insets of the panels in Figure 3C. On the horizontal axis, we plotted the number of factors (rank, *R*). The left panel shows results obtained for LFP responses when a cue stimulus was presented at angle *θ* = 0 degrees. The right panel shows similar results for a cue to *θ* = 60 degrees.

For both stimuli (and all other angles, Supplementary Figure 2A), the error reduced with an increasing number of factors (blue line in the insets). This provides a sanity check that the ALS algorithm produces meaningful results as the rank increased. This is similar to PCA, where the more principal components are included, the higher the variance explained. To find the optimal value for rank *R,* we computed two statistical measures: consistency and congruence.

Consistency was introduced as way to obtain the rank, *R*, in CD approximations in the paper ^48^. It uses certain elements of CD theory, known as CD factors, to compute an alternative approximation of the data matrix, known as Tucker3 approximation ^49^ and calculates the rank based on this approximation (see Methods for more details). High consistency suggests that there is a trilinear variation in the data. Congruence (also known as similarity) on the other hand, is the result of subtracting the maximum average uncorrected correlation coefficient (UCC) between factors corresponding to different initialisations of the ALS algorithm from 1 (see Methods). This addresses the local minima problem of the ALS algorithm. The lowest the congruence, the more stable the CD approximation (it does not depend on ALS initialisation). In these cases, congruence is small or close to zero, which was the case in our data too (see below).

To sum up, we chose the optimal rank *R* such that consistency is high and congruence is low. This ensures the CD approximation reflects a trilinear variation in the data and is stable. Consistency and congruence are shown in the right and left vertical axes of the bar plots in the main panels of Figure 3C. Consistency is shown using magenta bars, while congruence is shown using grey bars. Different bars correspond to different ranks. Rank is shown on the horizontal axes. Consistency values are shown on the left vertical axes, while congruence values are shown on the right vertical axes.

For cues presented at both *θ* = 0,60 degrees (left and right panels in Figure 3C) we obtained high consistency values for *R*=1,2 (magenta bars). The same was true also for other angles (Supplementary Figure 2A). For all values of *R* in Figure 3C and Supplementary Figure 2A, congruence was very small (grey bars). Its order was 10^-4^ for *θ* = 0 and 10^-5^ for *θ* = 60 degrees. Thus, in what follows, we used the CD approximation with *R*=2. For all angles, this corresponded to high consistency and low congruence. For *R*=2, the neuron mode was a matrix of dimensionality *N_S_* × 2. Fixing *m* = *M* *, where *M*\* = 1,2 we obtained two vectors 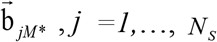 that approximate average LFPs across time and trials. These are the two *columns* of the neuron mode. Following ^47^, we call these vectors the 1^st^ and 2^nd^ neuron factors.

Recall that, each connectivity component is also a vector of length *N_S_*. In ^29^, we studied the first four connectivity components (similar to principal components in PCA). Here we focused on the first, as this explains most of the data variance similarly to the neuron factors that comprise the neuron mode. This explained about 35% of variance (Supplementary Figure 2B). Keeping up to 4 components, variance explained increased to about 60%. We asked whether the two neuron factors (recall *R*=2 above) were correlated with the first connectivity component. For cues presented at every angle (*θ* = 0,60, 120, 180, 240, 300 degrees), we computed the correlation coefficient and corresponding *p*-value between the 1^st^ and 2^nd^ neuron factors and the connectivity component averaged across trials. These are shown in Figure 3D. We found that the average connectivity component was significantly correlated with both the 1^st^ and 2^nd^ neuron factors. Correlations were significant for all angles. *P*-values (grey bars) are shown on the left vertical axes of left (1^st^ neuron factor) and right (2^nd^ neuron factor) panels. Only the 2 largest *p*-values for *θ* = 60 degrees are shown. All other *p*-values were much smaller than *p*< 10^-5^. Thus, they are not visible in the plot. The corresponding values of the correlation coefficient *r* are shown on the right vertical axes (burgundy bars). They are all high. Correlation coefficients were *r>*0.9 for all angles except *θ* = 60 for which *r*>0.7. Note that the CD approximation assumes a trilinear variation in the data, while the autoencoder approach we used to obtain our components is nonlinear. Thus, the remaining dissimilarity can be explained by a nonlinear mixing of latent states afforded by an autoencoder.

To sum up, we compared our approach for performing dimensionality reduction to a high dimensional SVD approach, known as Canonical Decomposition (CD^44,47^). We found that the effective connectivity component obtained using our approach correlates significantly with the neuron factors obtained using CD. This provides a second, independent validation of our approach.

All in all, we compared the effective connectivity components with results obtained using pairwise correlations and the latent states of a high dimensional SVD approach. Our components correlated with those found using alternative methods. Thus, all three methods found a similar structure of the latent space within which neural activity evolves, while neurons are maintaining cued angles.

### Stable electric fields emerge from neural ensembles that represent the same cued angle in different trials

To sum so far, we first found the latent space associated with maintenance of a cued angle (connectivity components). We then mapped this space to a cortical patch occupied by a neural ensemble—and obtained the connectivity kernels. These describe the exchange of information during cue maintenance. We found that the corresponding connectivity weights correlated significantly with single trial ensemble indices obtained using pairwise correlations ^40^ across a large percentage of trials.

Recall that the connectivity weights scale the input signal from other electrodes that targets a certain electrode measuring activity from a part of the neural ensemble. Having obtained these weights, we could then predict the Electric Field (EF) generated by the neural ensemble. The connectivity kernels describe how neurons communicate via electric signals sent from one part of the ensemble patch to the other. These electric signals generate the EF. Below we used the connectivity kernel and the deep neural field model to simulate EFs. We wanted to test if EFs were similar across trials where the same cued angle was maintained.

LFPs can be thought of as proxies to electric fields. However, it is not clear what their source is –and whether they are solely produced by neurons that participate in an ensemble or neighbouring neurons too. In other words, the ground truth regarding neural sources that produce the ensemble EF is unknown. Thus, we used our deep neural field model and the bidomain model to obtain predictions of ensemble EFs. The deep neural field model stands in for an in silico implementation of a neural ensemble. The bidomain model has been used to predict the electric field generated by biological tissues, like the cardiac muscle ^53,54^. To estimate the extracellular EF, the model requires only a measurement of the transmembrane potential *V^m^* (Methods). The bidomain model neglects ephaptic coupling and electromagnetic wave effects –that are small compared to electric effects. It yields the EF in the extracellular space by computing the Fourier transform of *V^m^* measurements and an analytical expression based on Bessel functions of the first and second kind.

Here, we obtained two EF estimates. First, EF estimates based on deep neural field model predictions of transmembrane potentials *V^m^*. These are simulated potentials after training the deep neural field model with all available data. We called them *simulated EFs.* Second, EF estimates based on real LFPs. These did *not* use the deep neural field model. LFPs were used as proxies for transmembrane potentials and replaced the simulated transmembrane potential from the neural field model above. We called the EFs obtained used real LFPs and the bidomain model, *real EFs*.

An example simulated EF estimate is shown in Figure 4A. The EF amplitude is shown on the vertical axis (*V/m*), while the two horizontal axes show the electrode number (ID; location on the cortex) and time (*ms*). *P*-values of correlations between EF amplitudes are shown in Figure 4B. These correspond to EFs generated by neural ensembles maintaining a cued angle at *θ* = 0 degrees. We here considered EF estimates from trials where our connectivity kernel correlated with the findings of ^40^ (correlated trials). To obtain these estimates, we first simulated neural activity using our deep neural field model. Variance explained was about 40% for all stimuli (cued angles, Supplementary Figure 5A and see also Figure 9 in ^29^; there we had used all trials, instead of correlated trials that we used here). After simulating neural activity, we computed EF estimates using the bidomain model. We asked whether they correlated across trials for the same electrode.

**Figure 4.**
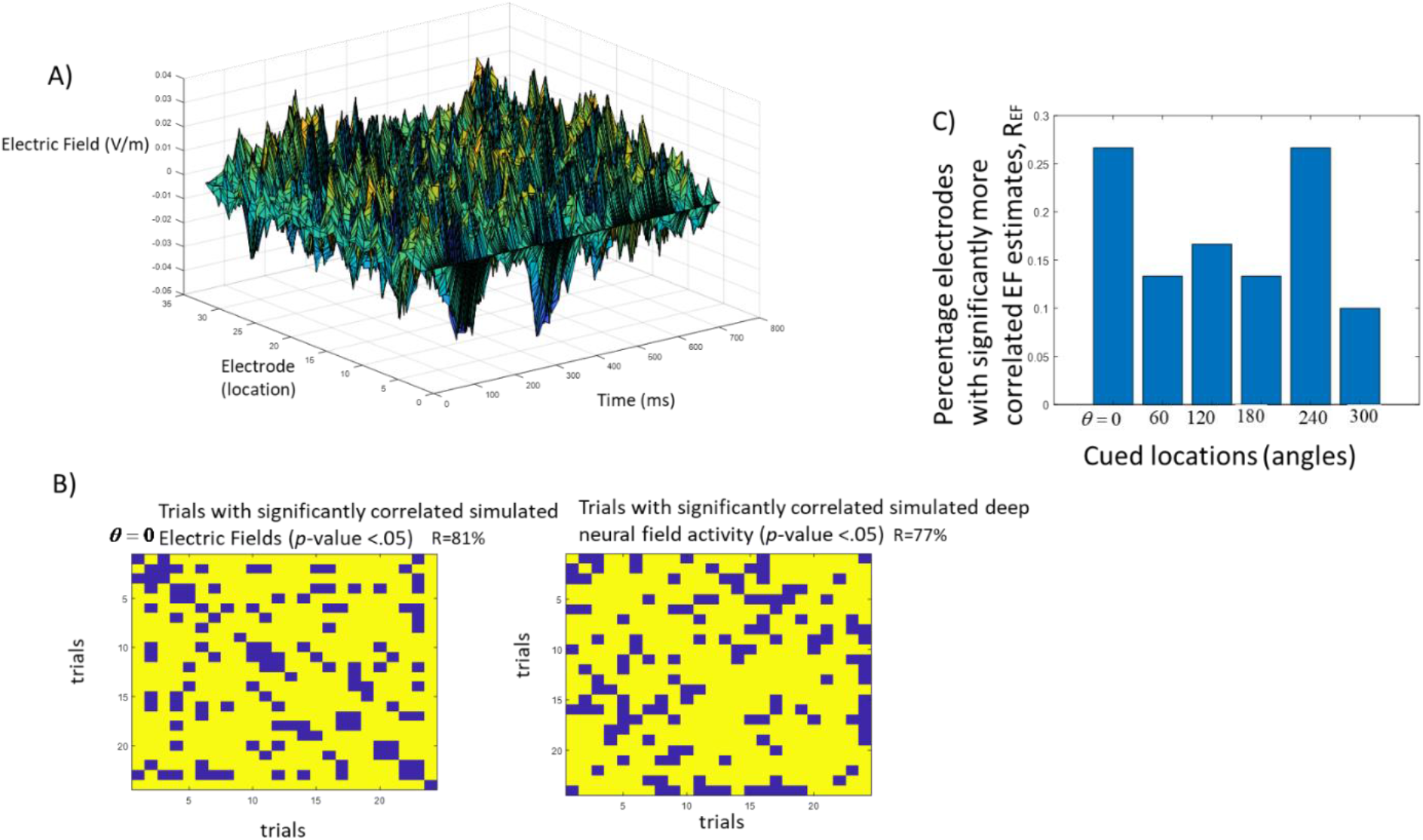
**A.** Example of simulated electric field (EF) using the bidomain model. The EF amplitude is shown on the vertical axis (*V/m*), while the two horizontal axes show the electrode (location on the cortex) and time (*ms*). **B.** (Left) *P*-values of correlations between single trial EF amplitudes. These correspond to EFs generated by neural ensembles maintaining a cued angle at *θ* = 0 degrees. Yellow entries in the correlation matrix denote significant *p*-values, *p* <.05. The percentage of significantly correlated single trial EF estimates is shown on the top right corner, *R*=80%. (Right) *P*-values of correlations between single trial deep neural field data. Yellow entries denote significant *p*-values as in Figure 4B. Percentage of correlated trials is lower than Figure 4B. **C.** Percentage of electrodes where electric field estimates were correlated across a larger number of trials compared to neural activity estimates, for different stimuli (angles).

Yellow entries in the correlation matrix denote significant *p*-values, *p <.05,* for an electrode at the edge of the patch and cued angle *θ* = 0. The percentage of significantly correlated single trial EF estimates is shown on the top right corner of the left panel in Figure 4B, *R*=81%.

Similarly, we found that EF amplitudes were also correlated across all trials and other angles with *R*= 70-80% (see Supplementary Figure 3). We also computed the corresponding correlation using deep neural field activity estimates at the same electrode for the same cued angle. The percentage of significantly correlated trials was *R*=77% (top right corner of right panel in Figure 4B). Note this is lower than the percentage of correlated trials computed using EF estimates obtained above.

We then asked if the same result holds across many electrodes: that is, if the percentage of correlated trials was lower when using single trial neural activity (deep neural field simulated data) compared to EF estimates. If it was, that would mean that neural activity was more variable than EF recordings. In other words, several distinct configurations of neural sources led to the same field.

To sum up, we asked whether for the same electrode (location on the ensemble patch), there was a significant difference in the percentage, *R,* of correlated trials obtained using: 1) reconstructed neural activity, which we called, *R_NA_* and 2) reconstructed EF estimates, called *R_EF_*. We repeated this for all electrodes and summarized results for each cued angle. Results are shown in Figure 4C. For all stimuli, a larger number of electrodes had reconstructed single trial EF estimates that were correlated across trials, *R_EF_*, compared to reconstructed neural activity estimates, *R_NA_*,:

Bars in Figure 4C show the percentage *electrodes, Q,* where *R_EF_* was significantly higher than *R_NA_*, *Q*=11-27%. To test for statistical significance, we used a Fischer exact test. This allows one to find differences between binomial distributions. Here, the binary variable describes whether a single trial EF estimate or neural activity estimate was correlated or not (the entries of matrices in Figure 4B). The null hypothesis was that there was no difference in the percentage of correlated trials (at the 5% significance level). We repeated the analysis for each electrode.

We found that a large part of electrodes had *R_EF_* > *R_NA_*^1^. In other words, electric field estimates were more often correlated across trials, i.e. more stable, compared to neural activity estimates^2^. In the next section, we will see that stronger stability of the EF compared to neural activity was also confirmed by decoding analyses. Training accuracy based on neural activity was significantly lower than accuracy based on EF estimates. This also suggests that information contained in delay neural activity was less stable than that contained in the electric field.

Not all electrodes had significant differences, *R_EF_* > *R_NA_*, because different stimuli activate different parts of the patch. Differences in connectivity components between stimuli are localised within those parts (see Figure 2 of ^29^ and relevant discussion). This suggests that only parts of the patch (certain electrodes) will be sensitive to changes of stimuli. EF and neural activity estimates measured at those electrodes will be correlated, that is, stable across trials (not the whole patch).

To sum, we found that electric fields were correlated across trials where the same cue was maintained. Further, the number of electrodes (locations) where this happens was larger than the corresponding number when neural activity was correlated. All in all, the above results suggest that stable electrical fields emerge from high-dimensional ever-shifting neuronal activity patterns of neural ensembles during trials where the same cue was maintained in memory networks. Having shown that electric fields are stable, we turned to the information carried by them and asked if it was stable too.

### Emergent electric fields carry unique information about working memory content

Finally, we asked whether EF produced by neural ensembles carried information about working memory content. We assessed whether EF estimates obtained using our approach were consistently different among neural ensembles that maintained different cued angles. In other words, we tested if we could distinguish between memorized cues based on EFs. If we could, this means that EFs can be uniquely associated with different working memories that are used to perform the task. To formally test this hypothesis, we used the EF estimates as classification features of different trials by cued angle. We used 450 trials and held out 20% of the data as a test set. We used simulated and real EFs as classification features and two different algorithms, Naïve Bayes and diagonal LDA. These are among the most commonly used.

The results of our analyses are shown in Figure 5 (using Naïve Bayes) and Supplementary Figure 7 (using diagonal LDA). Decoding accuracy values are shown on the vertical axis, while the corresponding electrodes (patch locations) are shown on the horizontal axis. We performed permutation tests, after shuffling class labels (cued angles) around. Blue bars denote observed accuracy values. Orange bars denote the maximum of the shuffled distribution. If blue bars are larger than orange, the observed accuracy is significantly higher than chance (max of shuffled estimates) at the *p*=0.01 level. This was the case for over half of the electrodes and accuracies obtained using simulated EFs (Figure 5A). The corresponding train and test confusion matrices are shown in Supplementary Figure 6A. These are averages over all electrodes. Accuracies were very similar for all stimuli^3^.

**Figure 5.**
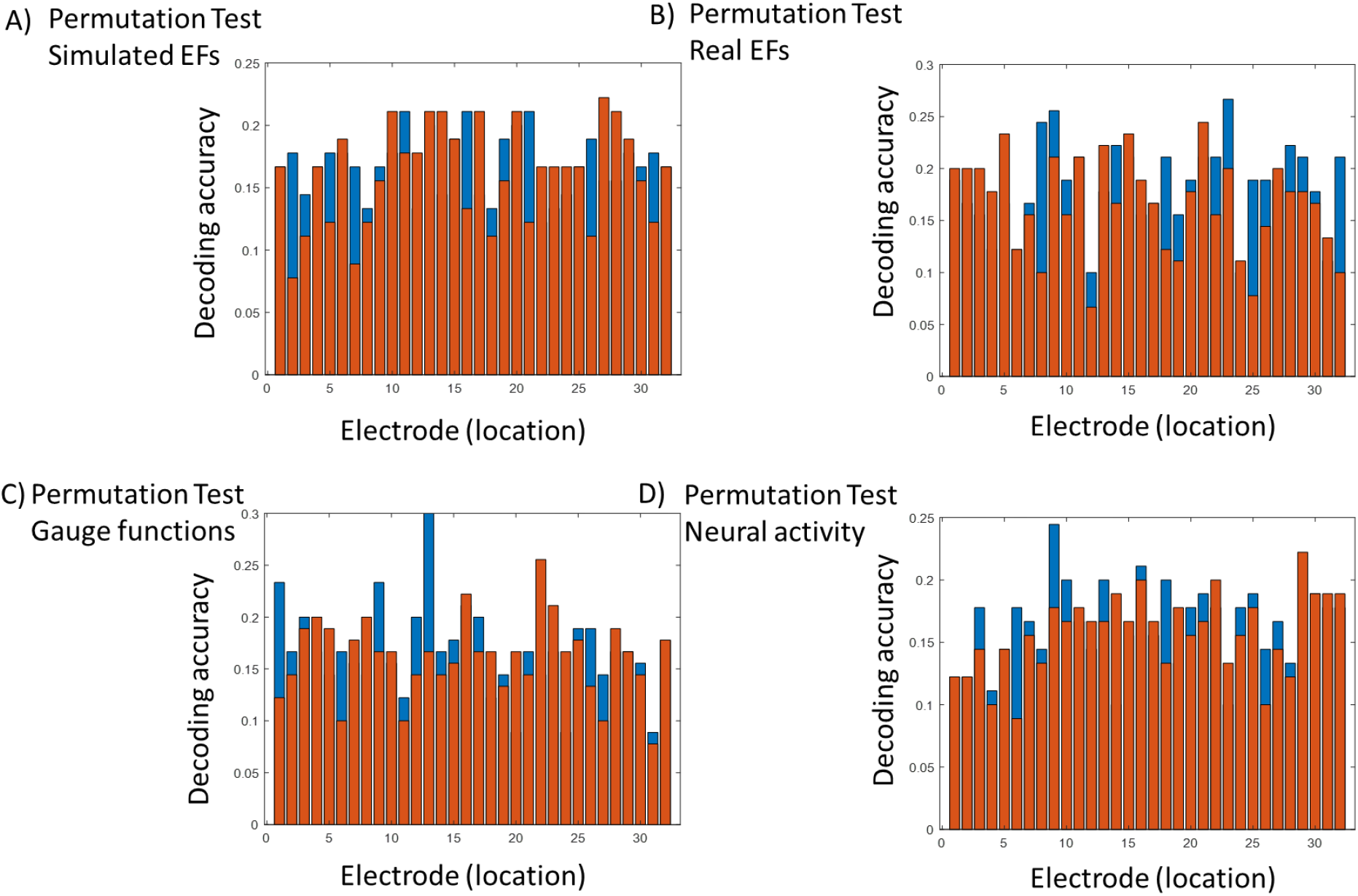
**A.** Permutation test of decoding accuracy based on simulated EF estimates. Accuracy values are shown on the vertical axes, while the corresponding electrodes (patch locations) are shown on the horizontal axes. Blue bars show observed accuracy estimates. Orange bars show the maximum obtained accuracy after performing *N*= 100 permutations. For those electrodes that unshuffled estimates are higher than the maximum of the distribution after shuffling, decoding accuracy is significantly higher than chance at the *p*=0.01 level. Over half of the electrodes have higher accuracy (blue bars) than the maximum of the distribution obtained after shuffling (orange bars). **B.** Same as in A. after replacing simulated EFs by real EFs. An equivalence test found that simulated and real EF accuracy estimate are the same (see text). **C.** Same as in A. after replacing simulated EFs by Gauge functions that do not depend on the neural field or dipole models. **D.** Same as in A. after replacing simulated EFs by neural activity estimates. A Welch test found that training accuracy obtained using neural activity was smaller than the corresponding accuracy obtained using real EFs (see text). Results shown in all panels were obtained using a Naïve Bayes classifier.

Recall that simulated EFs above were obtained from connectivity components, which, in turn, were obtained after training the neural field model on the whole dataset. Thus, decoding features contain some previous information from the data, something often referred to as data leakage. To address this, we computed the decoding accuracy using real EFs as features. Recall also that these were obtained after using LFPs as proxies for transmembrane potential. Thus, the corresponding accuracy will not be biased and includes out-of-sample validation based on a 20% held out test set. Similarly to simulated EFs, a permutation test confirmed accuracy significantly higher than chance at the *p*<0.01 level for over half of the electrodes (Figure 5B). The corresponding average confusion matrices are shown in Supplementary Figure 6B. Accuracies based on simulated EFs are similar to those obtained using LFPs (real EFs). To test for their equivalence, we used the TOST procedure^75^. We found that accuracies were the same, *t(31)=-2.05, p=*0.02 (assuming that a meaningful difference would be larger than 2%).

To sum up, we found that simulated and real EF estimates differed systematically depending on the exact cued angle; they were uniquely associated with the remembered stimulus. The above results confirm our earlier result that EFs were stable across trials where the same cued angle was maintained. They contained unique information about the remembered stimulus, that seems to be preserved across trials.

The theory of electromagnetism suggests that if EFs are stable, then the differences of the corresponding extracellular potentials should also be stable. These are known as *Gauge functions* (Methods). They are obtained by subtracting real LFPs recorded in different trials where the same cued angle was maintained. We thus asked if we could distinguish cued angles when using Gauge functions as decoding features. If we could, this would provide an alternative confirmation of our results. Crucially, Gauge functions do not rely on the validity of neither bidomain nor the deep neural field model. Thus, if they can distinguish between cued angles this is a confirmation of our result independent of these models.

The results of our analyses are shown in Figure 5C. As before, blue and orange bars correspond to observed accuracy and chance accuracy (maximum of the shuffled distribution) respectively. Accuracy obtained Gauge is similar to the results in Figures 5A and 5B. Thus, Gauge functions are also stable and contain information about the cued angle.

Finally, we repeated the decoding analyses using simulated neural activity (from the deep neural field model). Permutation test results are shown in Figure 5D. Accuracy was higher than chance (*p*<0.01) for over half of the electrodes. A one sided, *Welch* test also found that training accuracy based on neural activity was significantly smaller than accuracy obtained using real EFs *t(31*)=-8.2, *p*<0.001. The corresponding confusion matrices are shown in Supplementary Figure 6C. Correctly classified trials were fewer than those obtained using real and simulated EFs. Thus, neural activity did not contain the same stable information as the electric field. This is in accord with our earlier result (Figure 4).

All in all, we found that electric fields provided higher than chance decoding accuracy in predicting the remembered stimuli (cued angles). Thus EFs contained unique information about working memory content needed to perform the task.

## Discussion

We analyzed monkey LFP data from a spatial working memory task^28,29^. We found that stable electrical fields emerge from high-dimensional ever-shifting neuronal activity patterns of neural ensembles in the brain. We trained a biophysical neural network model as an autoencoder that learned to maintain spatial locations. This provided latent variables describing the connectivity of neural ensembles, which we called ‘connectivity components’. We also reconstructed single trial effective connectivity estimates, ‘ connectivity kernels’ ^29^. These describe information flow within the neural ensemble; in other words, the exchange of electric signals between neurons forming an ensemble. Crucially, this distinguishes our approach from other dimensionality reduction approaches ^5,7^. Our approach maps the latent space to a cortical patch. It goes beyond dimensionality reduction and reconstructs information flow.

Mathematically, the connection weights (kernel) can be thought of as the probability of having connections between neural populations forming a neural ensemble^29^. Other methods to obtain the probability function, including splines ^56^, and tools from complex systems ^58^. We will systematically consider these methods elsewhere. We here used a Restricted Maximum Likelihood (ReML) algorithm for obtaining the connectivity components^29^. This optimizes the same cost function used in variational autoencoders, called Free Energy (FE; also known as Evidence Lower Bound, ELBO). ReML does not require an explicit cross validation step (the E-step is embedded in the M-step after substituting the posterior variance). While cross validation (CV) partitions data in test and training sets, ReML prevents overfitting by penalizing for model complexity. The relationship between CV and FE for assessing source reconstruction error in the context of neuroimaging data has been systematically studied in several studies including^76,77^. CV error and FE are correlated^78^.

We found that the connectivity components and kernels were highly correlated with latent factors extracted by Canonical Decomposition (high dimensional SVD) ^44,47,50^. We also found that they were correlated with cluster indices obtained using unsupervised clustering ^40^.

Connectivity components and kernels describe the effective connectivity between different neurons forming a neural ensemble: How electric signals and information are exchanged between them. Using kernels and classic dipole theory of electromagnetism, we reconstructed the electric fields (EFs) produced by a neural ensemble. We reconstructed the electric field using three steps. Step 1 is obtaining the connectivity components after training a neural field model. These are the latent variables. Step 2 involves mapping the latent space to a cortical patch. This yields the connectivity kernel and predictions of neural activity. Step 3 is mapping neural activity to electric field using electromagnetism (bidomain model).

We found that different remembered locations resulted in different electric fields. These fields were highly stable across trials yet, at the level of specific circuits there was more variability (representational drift^17,18^). We reconstructed electric field and neural activity estimates during delay where the same location was remembered and looked at the percentage of trials where they were correlated. The percentage of electric field estimates that were significantly correlated across trials was higher than the corresponding percentage obtained using neural activity estimates and this was replicated across many electrodes and stimuli.

This result is also supported by the theory of electromagnetism. The same electric field can arise from different combinations of specific neurons and networks (electromagnetic sources and sinks^79^). This is known as non-uniqueness of the electromagnetic inverse problem: One cannot find the exact sources by measuring electric fields alone ^27^.

Across like trials, the same memory was maintained even though the inputs entering a given network changed. Electromagnetism predicts that neural sources will reconfigure themselves to accommodate these inputs but the overall electric field will be the same. When inputs change, the neural sources change but the electric field will not. This can explain of the observed variability in the patterns of neurons forming a neural ensemble. Here, we confirmed this hypothesis using LFP data and computational modeling. In future work, we will experimentally test the stability of the electric field.

Finally and importantly, different EFs were uniquely associated with different working memories needed to perform the experimental task successfully. To support this, it was shown that the EF estimates provided higher-than-chance decoding accuracies in predicting the remembered stimuli (i.e., cued angles). Further, training accuracy based on neural activity was lower and correctly classified trials were fewer than those obtained using EFs. Neural activity is less stable than the electric field.

Our model assumes that LFPs contain information about the excitation to inhibition (E/I) balance, despite being an aggregate measure of neural activity obtained from both excitatory and inhibitory populations. This is supported by both computational ^80–82^ and empirical ^83,84^ studies, see also ^85^ for a recent discussion. In particular, a large body of work by us and others using Dynamic Causal Models (DCM) has shown that it is possible to infer E/I ratios assuming that LFPs arise as a result of certain synaptic currents, usually AMPA and GABA_A_ currents, see e.g. ^86–91^. In future work, we will use separate recordings (depolarization or spike rates) from excitatory and inhibitory populations, to reconstruct excitatory and inhibitory activity separately.

In general, there are three different ways one can reduce the dimensionality in large, brain imaging datasets. Because these datasets involve three-way matrices (tensors) with dimensions (*time* x *neurons* x *trials*), three different sets of principal components (PCTs) can be obtained, in either (i) time; (ii) neurons (or channels) or (iii) trials domain. The outputs of this process are trajectories – i.e. collections of points– in domains spanned by the corresponding PCTs. For example, in ^73^ the output was motor timing (i.e. trajectories in a low dimensional domain spanned by time PCTs—ie. temporal evolution of population activity); while ^74^ obtained trajectories traced out by neurons in the motion and color domains (because PCTs along the second dimension, neurons, correspond to behaviourally relevant variables; neurons are grouped into PCTs depending on their tuning preferences). Finally, PCTs can be defined in the trial domain and the corresponding trajectories can then be used to obtain estimates of trial to trial variability. This can e.g. reveal changes in excitability of neural populations due to attention ^92^ and ongoing cognitive variables in general ^93^.

We here characterized the latent states during memory maintenance using biophysically informed models, neural fields. Because these models are defined in the time and neuron (i.e. space) domain, this reduction provides insights in both those domains. This, in turn, can help one understand the relation between representational drift and properties, like criticality^94,95^. Cortical dynamics in critical regimes are characterised by a co-occurrence of different temporal frequencies at different spatial scales ^96^. Both frequencies and spatial scales can be described by the connectivity components and principal axes obtained after training a neural field model^29^. In that earlier work, we showed that single trial principal axes predict the characteristic Lyapunov exponents that determine the timescales at which the system returns to equilibrium after perturbations, commonly known as critical slowing^97,98^. The connectivity components describe different neural ensembles, i.e. spatial patterns or combinations of neurons that maintain cued angles. These change between trials (representational drift). Thus, by studying single trial estimates of components and principal axes, one can link critical slowing with ensembles and representational drift. This will be considered elsewhere.

To sum up, we found that stable EFs emerge from high-dimensional ever-shifting neuronal activity patterns of neural ensembles in the brain. These EFs were robust across experimental trials where the same location was maintained, despite the continually changing neuronal activity, something known as the ‘representational drift’. Also, the low-dimensional emergent electrical fields carry information about working memories.

The stability of the electric field can allow the brain to control the latent variables (e.g., oscillations) that give rise to the same memory. We suggest that the electric field does not just emerge from the representational drift. It also helps sculpt and herd that general pattern of traffic. In other words, electric fields can act as “guard rails” that funnel the higher dimensional variable neural activity along stable lower-dimensional routes. We will test this hypothesis elsewhere. The low-dimensional stability in electric fields might help the brain perform computations, by allowing latent states to be reliably transferred between brain areas, in accord with modern engram theory ^99^. This is also in accord with the theory of Synergetics ^100–103^. The electric field can be viewed as a control variable similar to energy ^104^ and attention signals ^105^ that evolves more slowly than the latent variables that represent information. In other words, there might be a temporal hierarchy comprising the timescales of control parameters (e.g. electric field), order parameters (e.g. latent variables ^106,107^) and enslaved parts (e.g. oscillations/spiking ^102^).

All in all, our results and related work suggest that the electric field is conserved in memory networks and allows latent variables from different brain areas to interact and produce behavior. Although the exact neurons forming a neural ensemble differ from trial to trial (representational drift), the electric field is stable and contains unique information about the remembered stimulus, that seems to be preserved across trials.

## Acknowledgements

This work is supported by UKRI ES/T01279X/1, ONR MURI N00014-16-1-2832, and the JPB Foundation. We are also grateful to Dr Jay Myung, Jorge Chang and anonymous reviewers for useful comments.

## Supplementary Figures

**Supplementary Figure 1.**
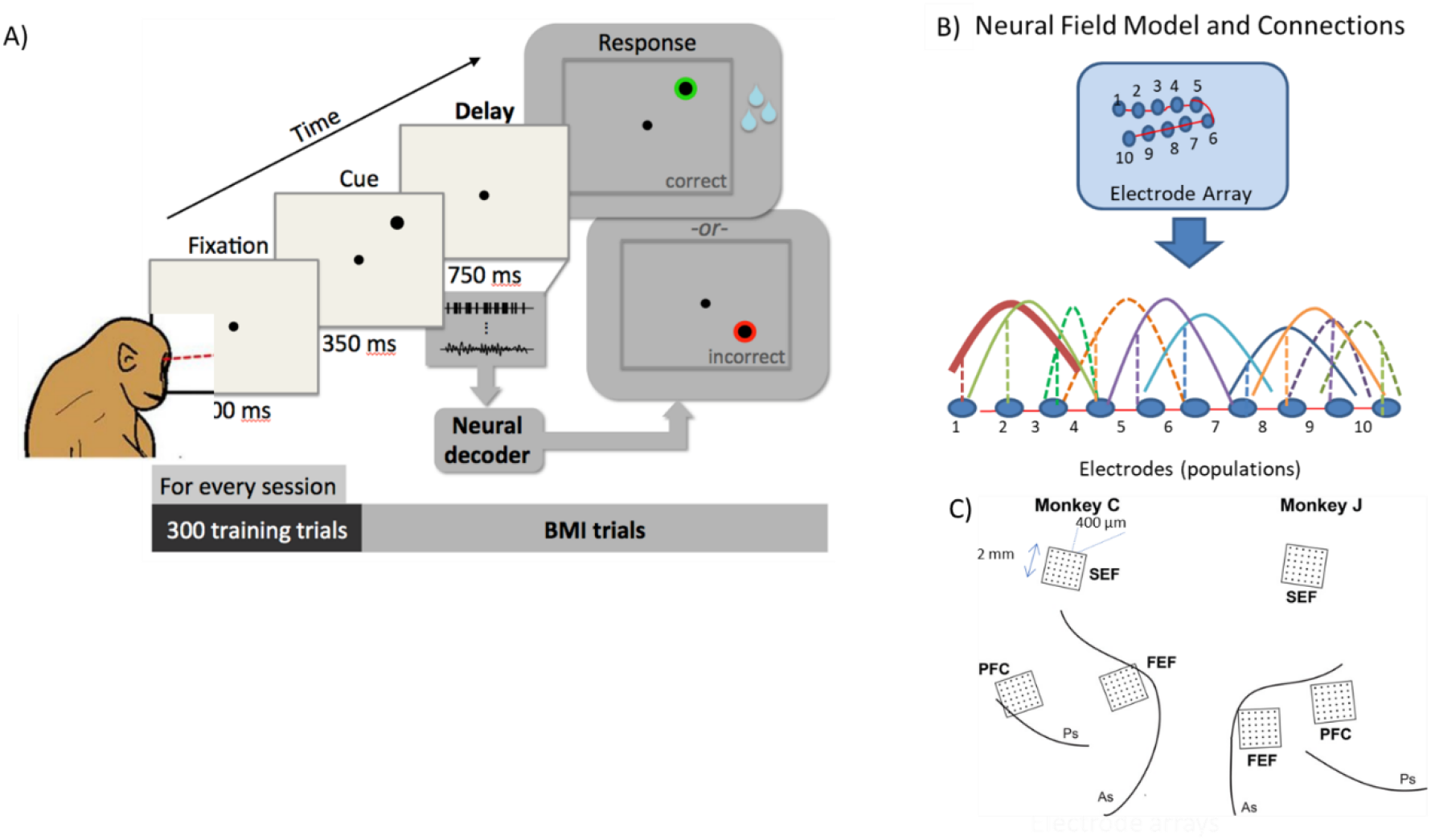
**A.** Oculomotor spatial delayed response task. Monkeys hold the location of one of six randomly chosen visual targets in memory over a brief delay period and then saccade to the remembered location. If a saccade was made to the cued angle, the target was presented with a green highlight and a water reward was delivered. Otherwise, the target was presented with a red highlight and reward was withheld. **B.** Deep neural field model and connections. This is a biophysical rate model. It is obtained as a simplification of a neural network model of coupled excitatory—inhibitory populations. It provides a quantitative way to describe each ensemble’s network interactions and patterns of activity across simultaneously recorded sites. The same model can describe different ensembles. Each electrode occupies a position on a cortical manifold *W* parameterized by the variable and is connected to all other electrodes with connections whose strength follows a Gaussian profile (coloured solid and dashed lines).**C.** 32-electrode chronic arrays were implanted unilaterally in PFC, SEF and FEF in each monkey. Each array consisted of a 2 x 2 mm square grid, where the spacing between electrodes was 400um.

**Supplementary Figure 2.**
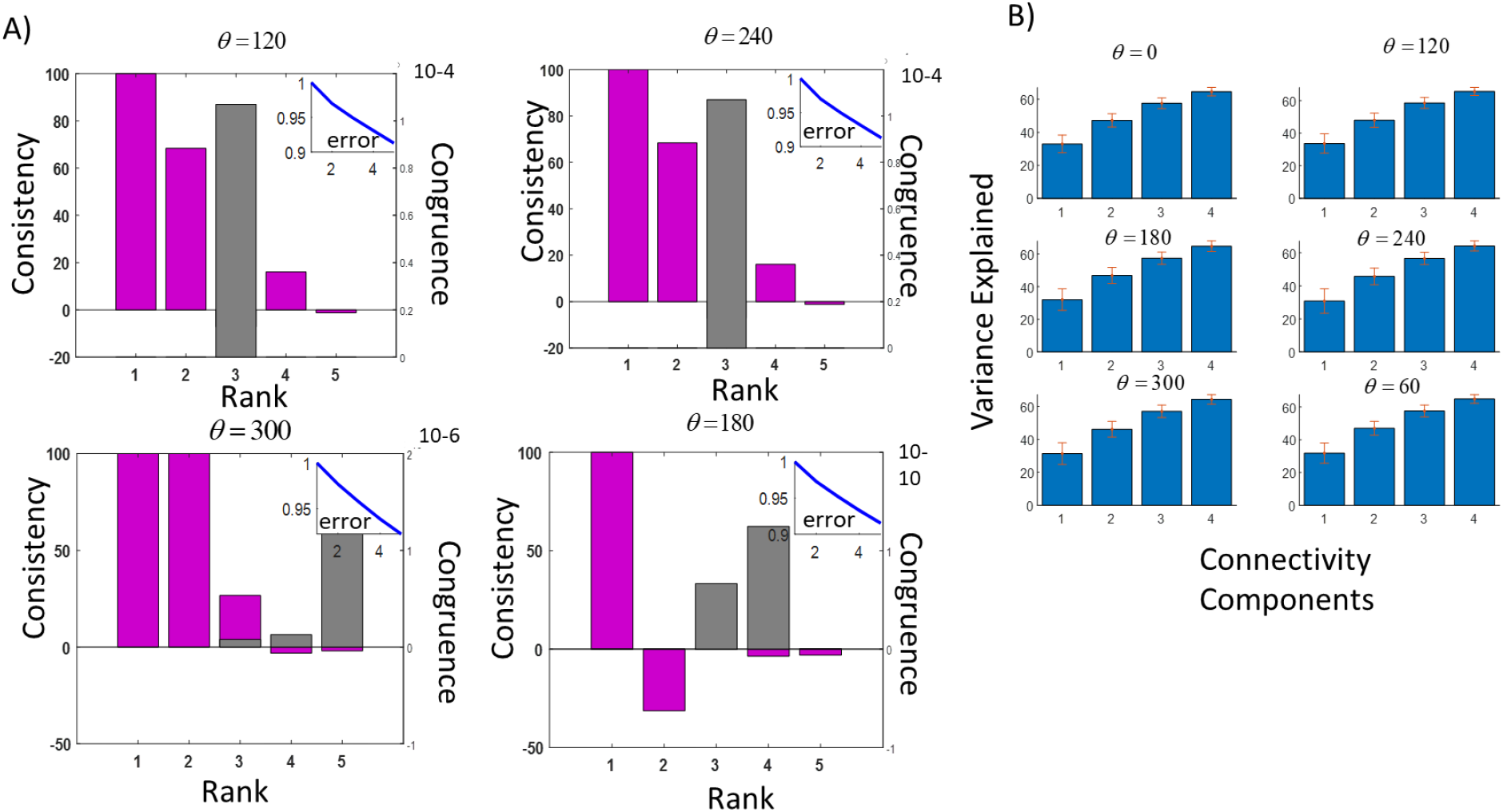
**A.** Results of Canonical Decomposition. Panels show consistency and congruence for cued angles at *0*=120, *θ*=240 (top) and *θ*=300, *θ*=180 (bottom) degrees. The format of each panel is the same as that of panels in Figure 1C. Consistency is shown using magenta bars, while congruence is shown using grey bars. **B.** Variance explained while keeping *d*=1,..,4 connectivity components.

**Supplementary Figure 3.**
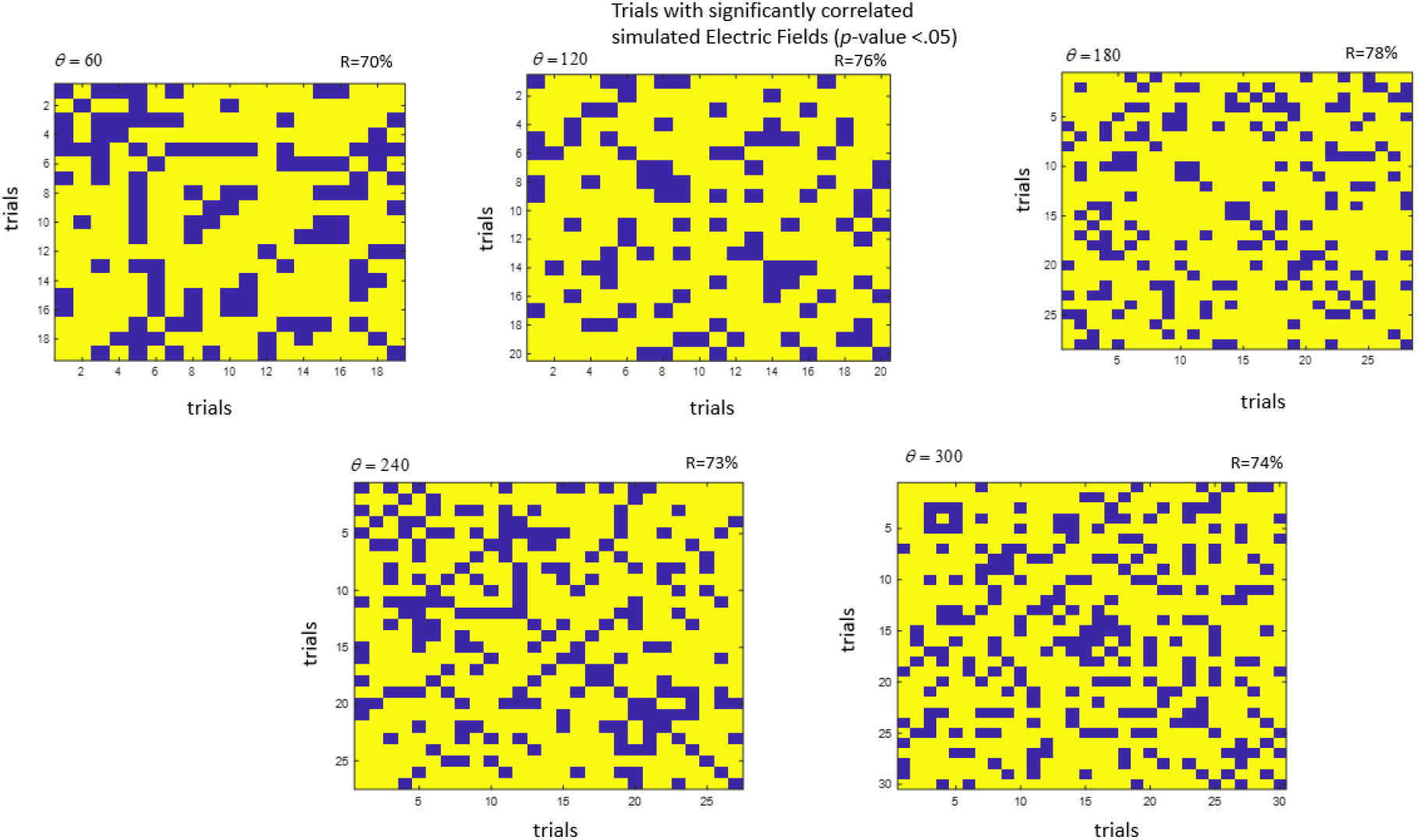
*P*-values of correlations between single trial EF amplitudes for cued angles at *0*=60, *θ*=120, *θ*=180 (top) and *0*=240, *θ*=300 (bottom) degrees. The format of each panel is the same as Figure 3B. Yellow entries in the correlation matrix denote significant *p*-values, *p*<.05.

**Supplementary Figure 4.**
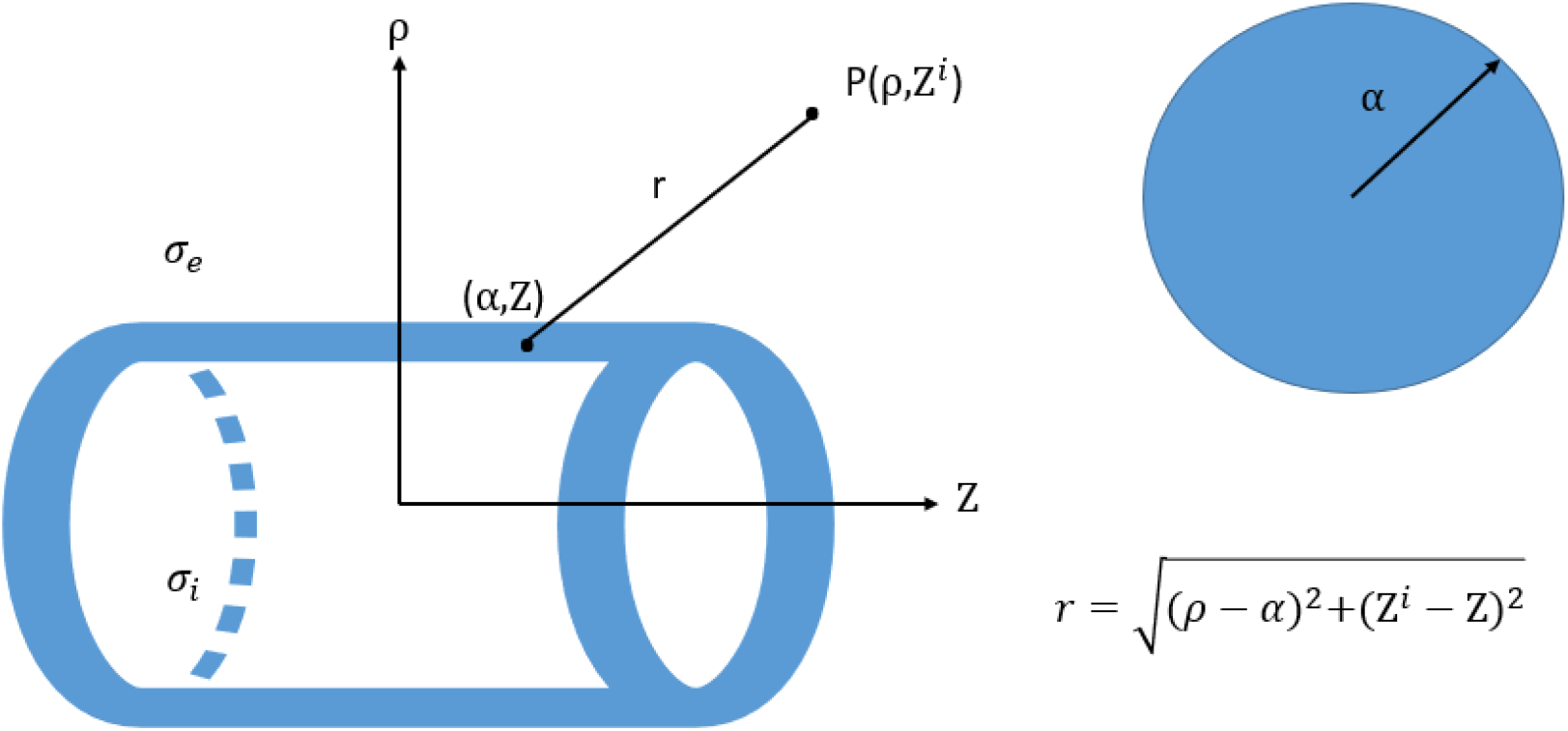
Bidomain model for the electric field generated by an active fiber in a semi-conductor (see Methods for the meaning of various symbols).

**Supplementary Figure 5.**
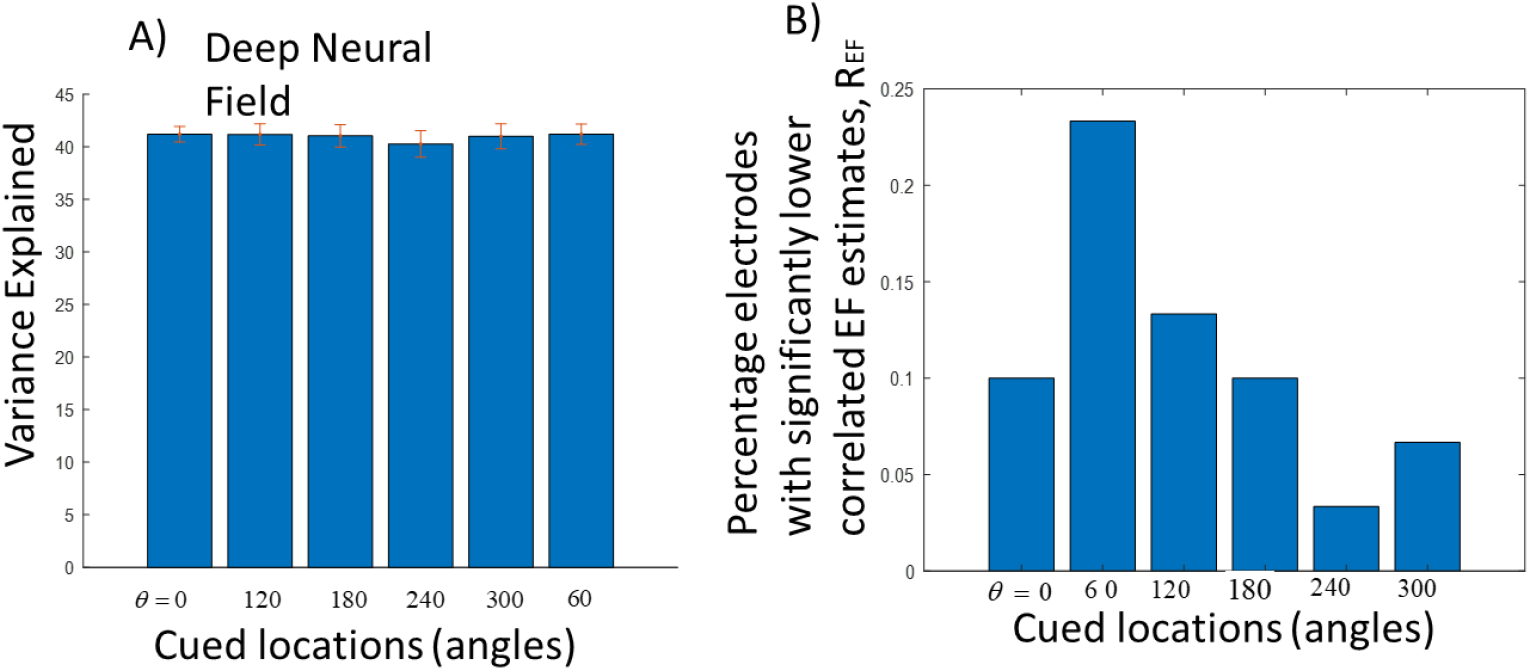
**A.** Variance explained after fitting the deep neural field model to real LFPs from different trials. Error bars denote standard deviation. Results are shown for different cued angles (angles). **B.** Percentage electrodes with significantly lower correlated EF estimates, R_EF_, for different cued angles.

**Supplementary Figure 6.**
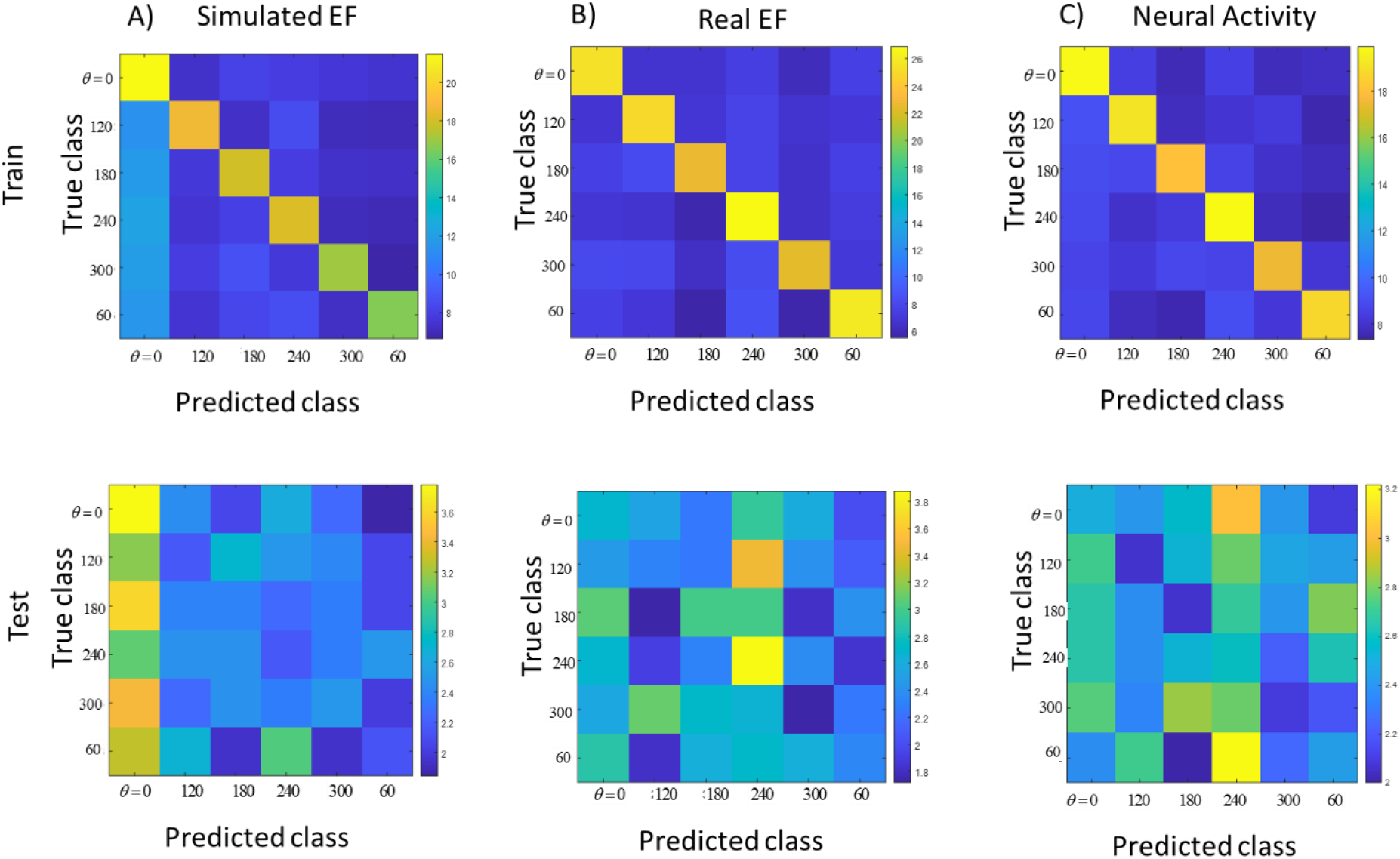
Breakdown of decoding accuracy through confusion matrices. Diagonal terms denote correctly classified trials. For each set of decoding features, we computed the corresponding train and test confusion matrices for each electrode (location on the patch). Then we averaged across electrodes. **A.** Train (top panel) and Test (bottom panel) confusion matrices, using Simulated EFs **B.** As in A. using Real EFs and **C.** Neural Activity (predictions of the deep neural field model).

**Supplementary Figure 7.**
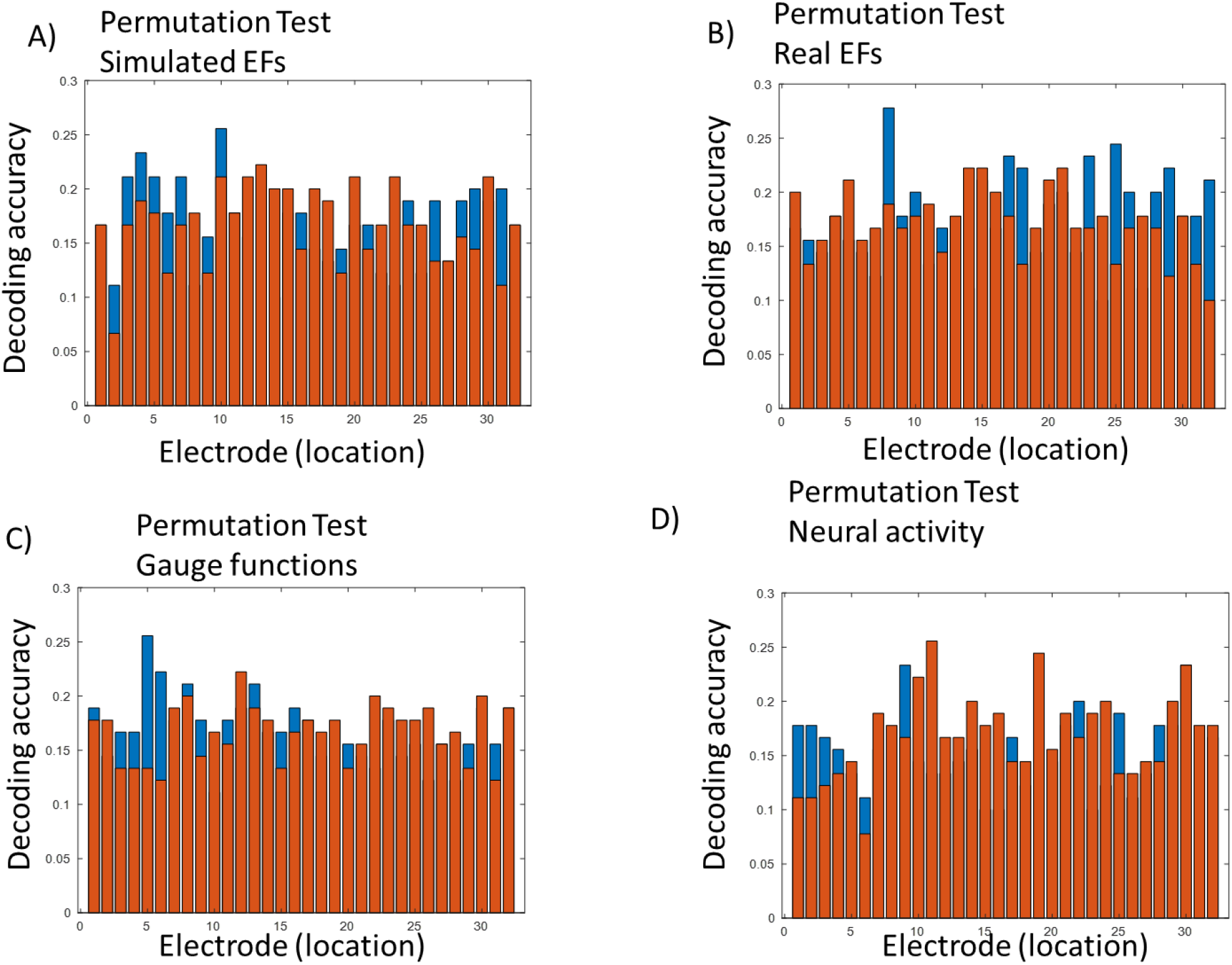
Permutation tests of decoding accuracy estimates obtained using diagonal LDA. The format of this Figure is exactly the same as the format of Figure 5.

## Footnotes

1 For all cued angles except *θ*=60 degrees (Supplementary Figure 5B).

2 During memory delay, some part of neural activity will be stable (attractor dynamics). This is not always picked up by EF estimates measured at certain locations (electrodes) due to assumptions in the bidomain model (isotropic field, homogeneous resistivity, infinite neural source etc; Supplementary Figure 5B).

3 Except for *θ*=0 degrees, which is slightly higher.

## Notes

### Competing Interest Statement

The authors have declared no competing interest.

